# An improved CRISPR-LICHT screen of 120 “Tdark” genes illuminates new biologically relevant SOX2 targets in human brain organoids

**DOI:** 10.64898/2026.10.23.761353

**Authors:** Giorgia Pozzolini, Simon Haendeler, Salma Amin, Roberta Baldi, Anna Cascio, Frank Edenhofer, Elena Taverna, Christopher Esk, Veronica Krenn, Silvia Nicolis

## Abstract

SOX2 is a transcription factor and central regulator of neural progenitor identity, yet the downstream molecular networks through which it supports human brain development remain incompletely characterized. Here, we applied CRISPR-lineage tracing at cellular resolution in heterogeneous tissue (CRISPR-LICHT) to human cerebral organoids to functionally interrogate SOX2-dependent regulatory networks. Using a SOX2-focused library targeting SOX2 together with conserved neurodevelopmental regulators expressed in neural progenitor cells, we showed that CRISPR-LICHT detects reproducible lineage-depletion phenotypes across genes already known to be involved in progenitor maintenance, fate specification, glial differentiation and neuronal maturation.

This extends the application of CRISPR-LICHT beyond the severe neural progenitor phenotypes previously characterized for microcephaly-associated genes and demonstrates its ability to detect perturbation effects across a broader set of neurodevelopmental regulators, involving SOX2 targets previously defined in mouse and now open to further understanding in a human model setting. We further screened 120 evolutionarily conserved, poorly characterized SOX2-dependent genes classified as “Tdark” within the “Illuminating the Druggable Genome framework” and identified 28 candidate hits whose disruption impaired neural progenitor lineage expansion and/or survival. These candidates included proteins associated with cytoskeletal organization, membrane dynamics and intracellular architecture, suggesting that SOX2-dependent developmental programmes may engage cellular processes beyond transcriptional regulation. Individual downregulation of CDC42EP4, a poorly characterized CDC42 effector linked to cytoskeletal organization, revealed altered distribution of targeted progenitors between ventricular- and subventricular-like compartments, with changes consistent with altered progenitor-state progression. Together, these findings expand the functional landscape of SOX2-dependent regulation in human neural progenitors and identify cellular architecture as a potential component of SOX2-associated developmental programs.

## Introduction

SOX2 is a transcriptional regulator crucial for brain development. Genetic evidence has established SOX2 as a dosage-sensitive regulator required for normal nervous system development across species. In humans, heterozygous pathogenic SOX2 variants cause SOX2 syndrome, a severe developmental disorder characterized by ocular abnormalities together with variable neurological manifestations, including brain malformations and intellectual disability (Ragge et al., 2005; Williamson et al., 2006; Kelberman et al., 2008; Williamson and FitzPatrick, 2014). In mice, complete loss of Sox2 causes early embryonic lethality, whereas conditional or hypomorphic mutations reveal stage- and region-specific requirements throughout the developing central nervous system (Avilion et al., 2003; Ferri et al., 2004; Favaro et al., 2009; Ferri et al., 2013; Mercurio et al., 2022).

SOX2 is primarily expressed in neural stem and progenitor cells and is widely used as a marker of neural stem cell (NSC) identity. Mechanistic studies, largely performed in mouse in vivo models or mouse NSC cultures (neurospheres), have shown that SOX2 regulates the balance between progenitor maintenance and differentiation, sustaining proliferation and self-renewal while preserving the competence to undergo lineage progression during neural development (Ferri et al., 2004; Favaro et al., 2009; Pevny and Nicolis, 2010; Mercurio et al., 2019; Mercurio et al., 2022; Marenco et al., 2026). These functions are context-dependent, with SOX2 dosage and developmental stage influencing its effects on neural progenitor maintenance and differentiation (Pevny and Nicolis, 2010). At the molecular level, SOX2 acts through extensive gene regulatory networks involving both promoter-proximal and distal regulatory elements. Genome-wide studies in mouse NSC cultures have shown that SOX2 preferentially occupies active distal enhancers engaged in long-range interactions with gene promoters and that Sox2 loss disrupts a substantial fraction of these interactions, accompanied by widespread changes in gene expression (Bertolini et al., 2019). Despite the large number of SOX2-regulated genes identified by genomic and transcriptomic analyses of mouse NSCs (Bertolini et al., 2019; Pagin et al., 2021), only a limited subset has been functionally characterized. These include Fos and Socs3, which contribute to NSC proliferation and long-term self-renewal (Bertolini et al., 2019; Pagin et al., 2021). Thus, although the transcriptional landscape downstream of SOX2 is increasingly defined, the functional contribution of many individual SOX2-regulated genes to NSC biology remains largely unknown.

Among the genes identified through transcriptomic analyses of Sox2-deficient mouse NSCs is a subset of evolutionarily conserved, poorly characterized genes encoding proteins classified as “Tdark” within the Illuminating the Druggable Genome framework (Nguyen et al., 2017; Oprea et al., 2018; Bertolini et al., 2019). This category encompasses proteins for which limited functional information is available. Functionally interrogating these understudied SOX2-regulated genes therefore provides an opportunity to move from transcriptional association to functional characterization and to identify previously unrecognized components of SOX2-dependent developmental programmes.

While much of our mechanistic understanding of SOX2 function derives from mouse models, studies in human neural systems indicate that SOX2 similarly occupies a central, but highly context-dependent, position in neural progenitor regulatory networks. In human embryonic stem cell-derived neural progenitor cell cultures (NPCs), SOX2 perturbation affects progenitor proliferation and neuronal differentiation, with the SOX2-LIN28/let-7 axis providing one mechanism linking SOX2 activity to progenitor maintenance and neurogenic competence (Cimadamore et al., 2013). Genome-wide analyses further showed that SOX2 engages distinct cell-state-dependent transcriptional programmes during the transition from pluripotency to neural progenitor identity (Zhou et al., 2016). More recently, single-cell CRISPR interference in primary human cortical radial glia identified SOX2 as part of a complex transcription factor network controlling radial glial state and lineage composition during human cortical neurogenesis (Ding et al., 2026). Importantly, the expression of SOX2 in neural progenitors is also recapitulated in human cerebral organoids, where SOX2- positive progenitors populate ventricular zone-like regions that reproduce key features of the developing human brain. These three-dimensional models provide an opportunity to investigate neural progenitor regulatory mechanisms in a developmental context that recapitulates progenitor expansion, neuroepithelial and radial glial organization, and neuronal lineage progression (Lancaster et al., 2013; Velasco et al., 2019). Together, these studies indicate that SOX2 has conserved functions in human neural progenitors while also highlighting the complexity and cell-state dependence of its regulatory network.

Advances in pooled CRISPR screening, combined with human brain organoid models, have opened new opportunities to functionally interrogate genes involved in human neurodevelopment in a three-dimensional context. Such approaches enable systematic analysis of genes controlling neural progenitor fitness, expansion and developmental progression (Esk et al., 2020; Li et al., 2023; Meng et al., 2023). In particular, CRISPR- lineage tracing at cellular resolution in heterogeneous tissue (CRISPR-LICHT) provides a lineage-resolved strategy to identify genetic perturbations affecting NSC expansion in cerebral organoids and has previously been used to interrogate genes associated with microcephaly, a neurodevelopmental condition characterized by reduced brain size (Esk et al., 2020). Given the central role of SOX2 in NSC biology, this approach provides a framework to systematically investigate the functional contribution of genes within the SOX2-regulated network in human neural progenitors.

Here, we applied CRISPR-LICHT to identify regulators of NSC abundance within the SOX2- regulated gene network. First, using a SOX2-focused library targeting SOX2 alongside conserved neurodevelopmental regulators, we benchmarked and streamlined the lineage-resolved screening workflow for the detection of perturbations affecting neural progenitor abundance. We then applied this framework to a library targeting 120 evolutionarily conserved, poorly characterized SOX2-dependent genes classified as Tdark and identified 28 candidate regulators of NPC abundance linked to cytoskeletal organization, membrane dynamics and intracellular architecture. Among these, functional characterization of CDC42 effector protein 4 (CDC42EP4), a CDC42-binding protein implicated in cytoskeletal organization, revealed a requirement for appropriate neural progenitor lineage progression, providing a functional link between the SOX2-regulated network and cytoskeletal control. Together, our findings expand the functional landscape of SOX2-dependent regulation in human neural progenitors and establish CRISPR-LICHT as an effective approach for functional interrogation of poorly characterized components of developmental gene-regulatory networks.

## Results

### Design of complementary CRISPR-LICHT libraries to interrogate SOX2-dependent regulators of early brain development

We previously identified approximately 900 genes downregulated upon Sox2 loss in murine primary neural stem cells using transcriptomic analysis (Bertolini et al., 2019). These included genes with established roles in murine neurodevelopment (such as Egr1, Zeb2 and Gria2), as well as genes with limited functional annotation. Among these approximately 900 downregulated genes, 120 encoded poorly characterized proteins classified as Tdark according to Pharos at the time of library design and were conserved in humans (as defined by the presence of a corresponding human homolog), revealing a substantial set of evolutionarily conserved but functionally unexplored targets downstream of Sox2.

To move from transcriptional association to functional testing in a human model of early brain development, we applied CRISPR-LICHT, a screening strategy in cerebral organoids that combines CRISPR-Cas9 mutagenesis with lentiviral lineage barcoding to identify perturbations affecting NPC abundance across large sets of candidate genes (Esk et al., 2020). In this system, each genetic perturbation is linked to a clonal barcode, allowing lineage size to be used as a quantitative readout of progenitor expansion and survival during organoid development.

We designed two complementary CRISPR-LICHT-compatible libraries to interrogate the SOX2-dependent regulatory network at different levels (Fig. 1A). To determine whether CRISPR-LICHT could detect changes in NPC abundance caused by perturbation of SOX2 and established SOX2-dependent neurodevelopmental regulators, we designed a targeted reference screen using a SOX2-focused library. This library included gRNAs targeting *SOX2* itself together with 12 SOX2-regulated genes identified in Bertolini et al. (2019), namely *ZEB2, KCNA2, GJA1, FBRSL1, EGR1, EGR2, GRIA2, PPP2R2B, CNTN1, GADD45G, KCNK2* and *ARX*, for which neurodevelopmental functions have previously been reported (Table S2). Protein-class and Gene Ontology analyses showed that these genes encode primarily transcription factors and chromatin regulators, together with ion channels, signalling proteins and structural or cytoskeletal components (Tables S2-S4). Consistently, they are associated with a broad range of neurodevelopmental processes, including transcriptional regulation, progenitor maintenance, fate specification, glial differentiation, neuronal maturation, excitability and axonal organization (Fig. 1B,C). Mutations in several of these genes are also associated with human neurological disorders, including epilepsy (*ZEB2, KCNA2, GRIA2, KCNK2*), intellectual disability (*ZEB2, EGR1, ARX*) and brain malformations (*ZEB2, PPP2R2B*) (Table S2). Their documented roles in neural development therefore provided a reference set against which the ability of the screening workflow to recover perturbation-associated changes in NPC abundance could be assessed.

**Figure 1.**
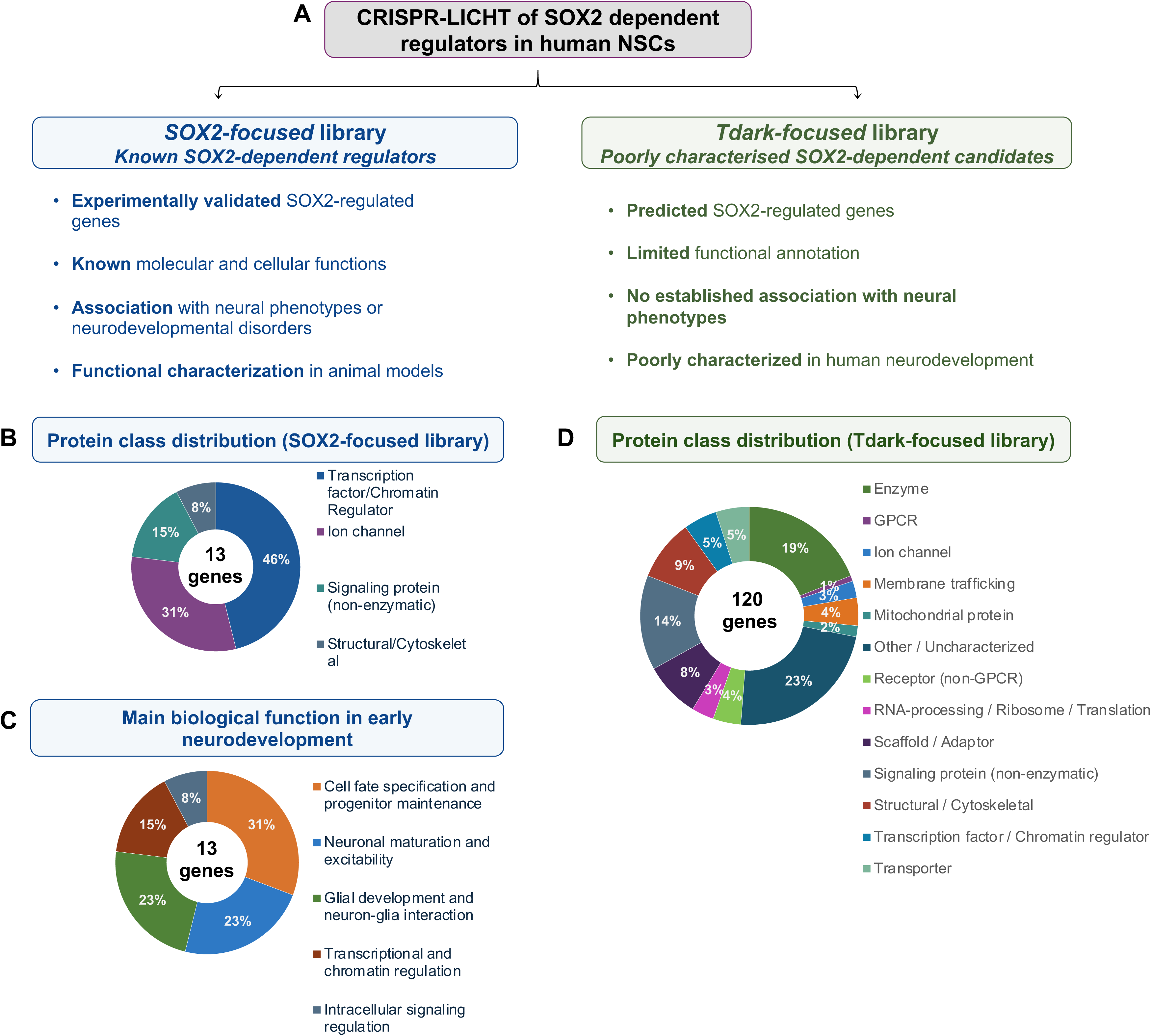
Design and characterization of CRISPR-LICHT libraries targeting SOX2-dependent regulators in human NSCs **(A)** Schematic overview of the two custom CRISPR-LICHT libraries generated to investigate SOX2-dependent regulators in human neural stem cells (NSCs). The *SOX2-focused library* includes experimentally validated SOX2-regulated genes with established molecular and cellular functions, reported associations with neural phenotypes or neurodevelopmental disorders, and prior functional characterization in animal models. The *Tdark-focused library* comprises poorly characterized SOX2-dependent candidate genes predicted to be regulated by SOX2, with limited functional annotation and no established association with neural phenotypes or human neurodevelopment. **(B)** Protein class distribution of genes included in the SOX2-focused library. Genes were classified according to their annotated molecular functions and protein families, including transcription factors/chromatin regulators, ion channels, signaling proteins, and structural/cytoskeletal components. **(C)** Main biological processes associated with genes included in the SOX2-focused library during early neurodevelopment. Functional categories include cell fate specification and progenitor maintenance, neuronal maturation and excitability, glial development and neuron-glia interaction, transcriptional/chromatin regulation, and intracellular signaling pathways. **(D)** Protein class distribution of genes included in the Tdark-focused library. Genes were grouped based on predicted or annotated protein classes, including enzymes, GPCRs, ion channels, membrane trafficking proteins, mitochondrial proteins, receptors, scaffolding/adaptor proteins, signaling proteins, structural/cytoskeletal proteins, transcription/chromatin regulators, and transporters. Notably, a significant proportion of candidates corresponded to poorly characterized or unannotated proteins, highlighting the limited functional knowledge currently available for Tdark genes.

To functionally interrogate poorly characterized genes within the SOX2-dependent network, we additionally designed a discovery library targeting 120 Tdark genes (Fig. 1A,D; Bertolini et al., 2019). These genes were selected from the subset of transcripts downregulated following *Sox2* loss in murine neural stem cells that were evolutionarily conserved in humans, and also classified within the Tdark category of the Illuminating the Druggable Genome framework at the time of library design. We therefore refer to this library as the Tdark-focused library. In contrast to the SOX2-focused library, the Tdark-focused library displayed a broader and more heterogeneous protein-class composition, including enzymes, signalling proteins, structural or cytoskeletal proteins, scaffold or adaptor proteins, transporters, receptors, membrane-trafficking proteins and mitochondrial proteins (Fig. 1D). A substantial fraction (23%) comprised largely uncharacterized proteins, further highlighting the limited functional annotation of the targeted gene set. The library also included 13 positive controls and 13 non-targeting controls. Positive-control gRNAs were designed in the original CRISPR-LICHT study to target genes whose disruption was expected to induce NPC depletion, whe reas non-targeting controls were not expected to affect NPC abundance (Esk et al., 2020) (Tables S 5, S6).

### Establishment of the CRISPR-LICHT culture setting

To implement CRISPR-LICHT, we first assessed whether the inducible eCas9-dTomato knock-in line used in the original CRISPR-LICHT study could generate brain organoids under our culture conditions. This line carries an eCas9-dTomato cassette inserted into the AAVS1 safe-harbour locus, which is activated by Cre-ERT2-mediated recombination following 4-hydroxytamoxifen (4-OHT) treatment (Esk et al., 2020) (Fig. 2A). We differentiated these cells into brain organoids using an adapted version of the Lancaster et al. protocol (Lancaster et al., 2013) (Fig. S1). Morphological inspection up to day 30 and immunostaining confirmed organoid growth and the presence of PAX6- and FOXG1-positive cells organized into neuroepithelial structures, hereafter referred to as ventricular zone-like (VZ-like) regions, consistent with the emergence of forebrain-like progenitor domains (Fig. 2A,B). We also detected robust SOX2 expression within VZ-like structures, confirming the presence of SOX2-positive neural progenitor populations (Fig. 2C; Fig. S1).

**Figure 2.**
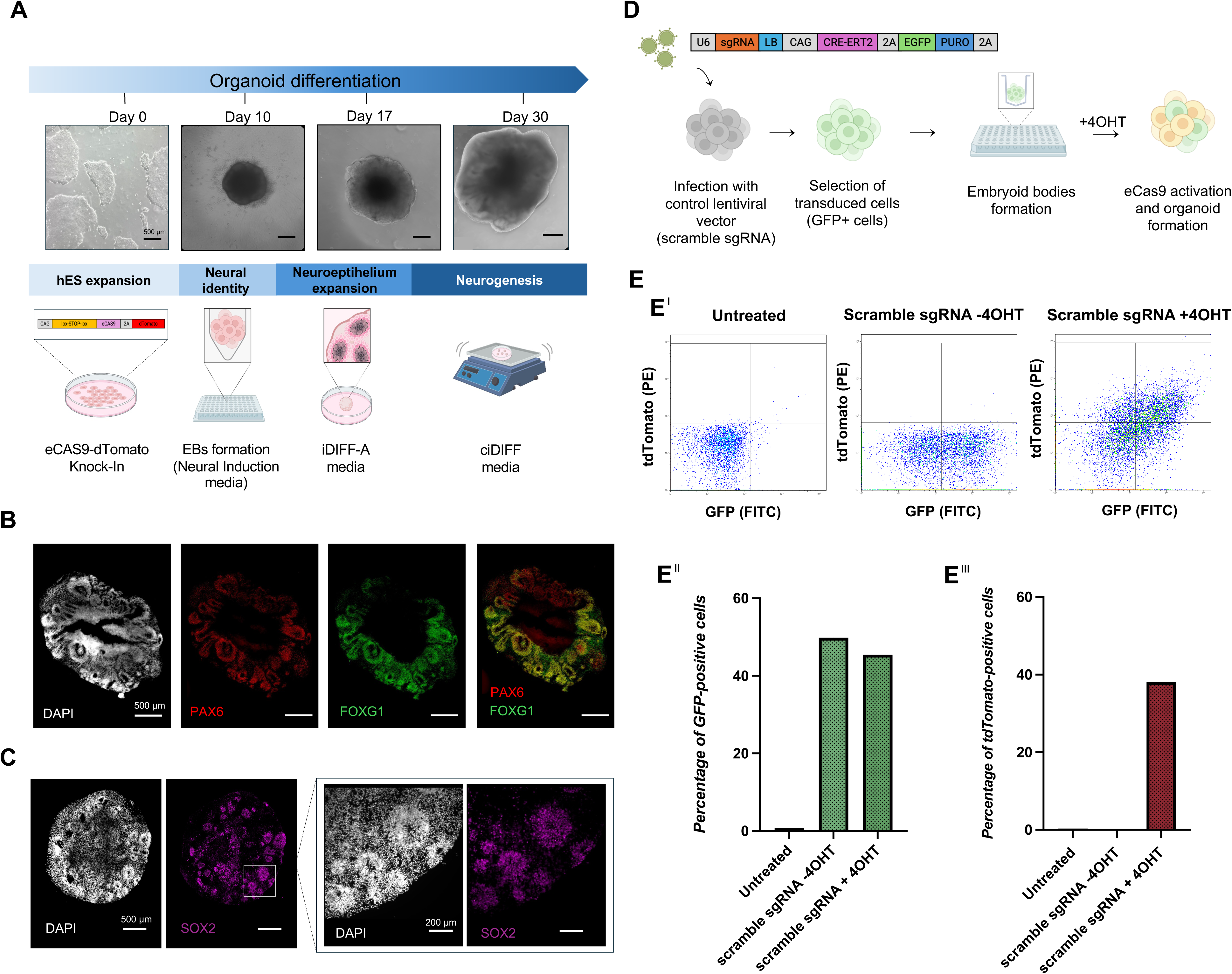
Establishment and validation of the CRISPR-LICHT platform in human cortical organoids **(A)** Experimental workflow for generation of inducible CRISPR-LICHT human cortical organoids using the H9-CAS9 cell line. The H9-CAS9 line carries an inducible enhanced Cas9 (eCas9) fused to a dTomato reporter under the control of a lox-STOP-lox cassette inserted into the AAVS1 safe harbour locus. Following lentiviral delivery of Cre-ERT2, transduced cells were differentiated into cortical organoids using a modified Lancaster protocol. Cas9 activation was induced by 4-hydroxytamoxifen (4-OHT) treatment during differentiation, enabling temporal control of genome editing. Representative brightfield images show organoid morphology at different stages of differentiation (days 0, 10, 17, and 30). **(B)** Immunofluorescence analysis of day 30 cortical organoids showing expression of dorsal forebrain progenitor markers PAX6 and FOXG1. The spatial organization and marker distribution are consistent with neuroepithelial structures characteristic of developing cortical tissue. Scale bars, 500 µm. **(C)** Representative immunofluorescence images showing robust SOX2 expression within neural rosette-like regions of day 30 cortical organoids. Higher magnification images highlight SOX2-positive neuroepithelial domains consistent with neural progenitor identity. Scale bars, 500 µm (left panels) and 200 µm (right panels). **(D)** Schematic overview of the pilot validation experiment used to assess lentiviral transduction efficiency and inducible Cas9 activation independently of the CRISPR-LICHT gRNA libraries. H9-CAS9 cells were transduced with a control lentiviral vector carrying the CRISPR-LICHT backbone and a scramble sgRNA, followed by enrichment of GFP- positive cells, embryoid body formation, and cortical organoid differentiation. Cas9 activation was induced by 4-OHT treatment during differentiation. **(E)** Flow cytometry analysis of dissociated cortical organoids between days 30 and 35 following scramble sgRNA transduction and inducible Cas9 activation. Organoids without 4-OHT treatment retained predominantly GFP-positive/dTomato-negative cells, whereas 4-OHT-treated organoids displayed a distinct dTomato-positive population, confirming efficient Cre-ERT2-dependent activation of the inducible eCas9 cassette. Quantification of GFP and dTomato populations is shown in the accompanying bar plots.

We next assessed lentiviral transduction and inducible eCas9 activation independently of the screening libraries using a control CRISPR-LICHT vector carrying a scramble sgRNA (Fig. 2D; Fig. S2). Following lentiviral transduction and enrichment of GFP-positive pluripotent stem cells by fluorescence-activated cell sorting (FACS), sorted cells were aggregated to form organoids and were treated with 4-OHT at day 5 of differentiation. Flow cytometry analysis of dissociated organoids between days 30 and 35 was used to assess eCas9-dTomato activation. Untreated organoids remained predominantly GFP-positive and dTomato-negative, whereas 4-OHT-treated organoids contained a dTomato-positive population, confirming Cre-ERT2-dependent activation of the eCas9-dTomato cassette (Fig. 2E). Although organoids were generated from GFP-positive transduced cells and dTomato expression could therefore only be induced in transduced cells, a fraction of GFP-negative cells was detected at the endpoint of the experiment within the dTomato-positive population. This observation is consistent with silencing of the GFP-expression cassette in a subset of transduced cells during organoid differentiation.

Together, these data established that the inducible eCas9-dTomato line supports brain organoid generation under our culture conditions, maintains SOX2-positive neural progenitor populations and permits detectable activation of the eCas9-dTomato cassette at the screening endpoint.

### CRISPR-LICHT screening designs for SOX2- and Tdark-focused libraries

We performed CRISPR-LICHT screens with the SOX2-focused and Tdark-focused libraries in parallel, using 30–35 days in vitro (div) as the endpoint for analysis. This window was selected to interrogate perturbation effects during a SOX2-positive, progenitor-enriched stage, as supported by the robust SOX2 expression observed at day 30 (Fig. 2C). In both screens, each gRNA was coupled to a lineage barcode, enabling simultaneous identification of the genetic perturbation and quantitative tracking of the corresponding lineage during organoid differentiation. Because lineage size integrates the proliferative capacity and survival of targeted progenitors over time, barcode abundance was used as a quantitative readout of lineage expansion and relative NPC abundance within heterogeneous organoid tissue.

Both libraries were initially designed around the standard CRISPR-LICHT workflow described by Esk et al. (2020), which relies on FACS enrichment of GFP-positive transduced pluripotent stem cells before organoid generation, induction of eCas9 activity by 4-OHT during organoid differentiation, and endpoint FACS isolation of dTomato-positive cells. However, the extent to which the initial enrichment and endpoint isolation steps influence screening performance had not been systematically assessed. We therefore used the lower-complexity SOX2-focused library, which contained established neurodevelopmental regulators with expected perturbation-associated phenotypes, to compare alternative processing strategies. Specifically, we tested two technical variables: puromycin selection as an alternative to GFP-based FACS enrichment of transduced cells before organoid generation, and bulk harvest following organoid dissociation as an alternative to endpoint FACS isolation of dTomato-positive cells (Fig. 3A).

**Figure 3.**
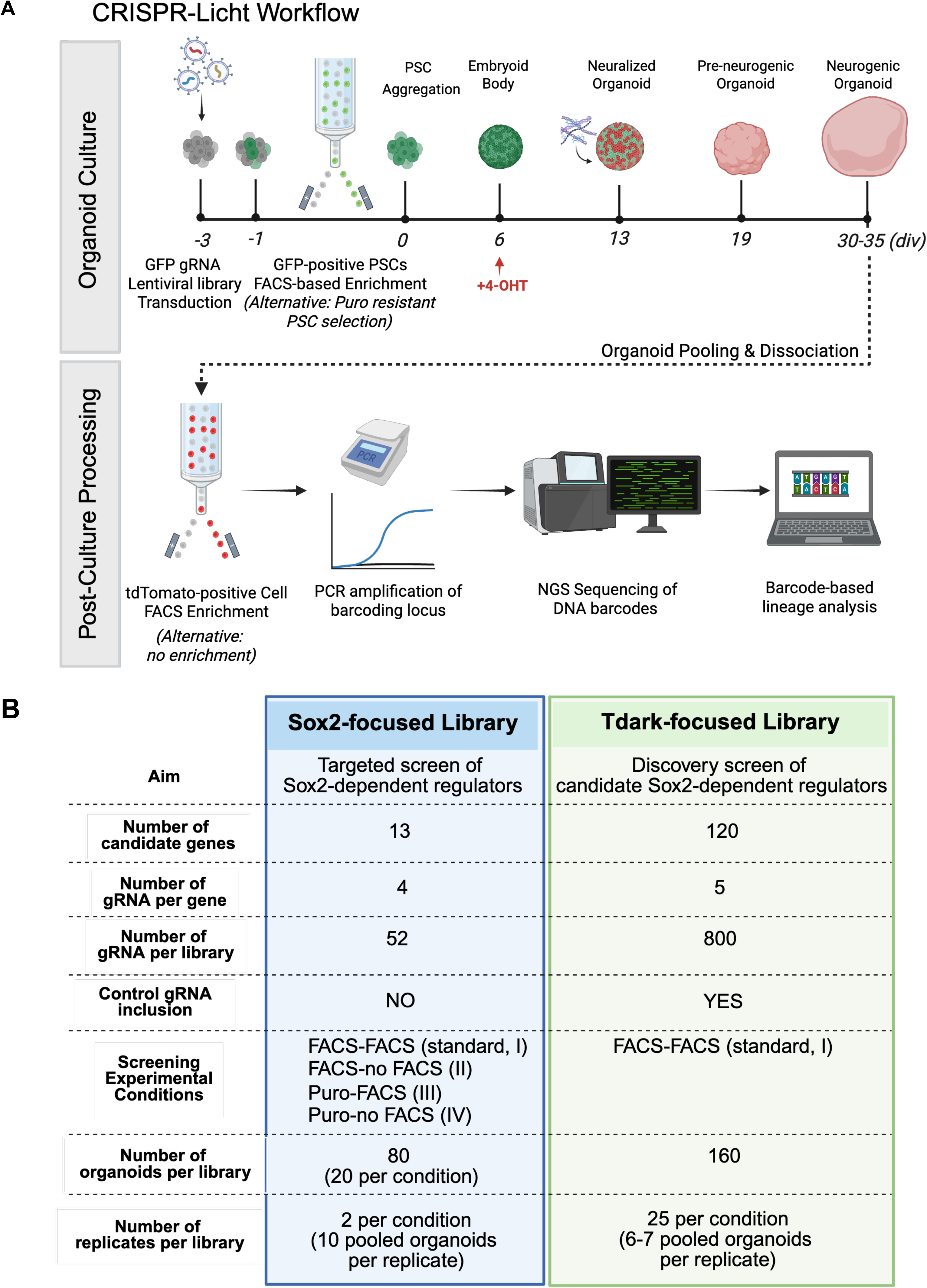
Experimental workflow and design of SOX2- and Tdark-focused lineage-resolved CRISPR screens in human cerebral organoids. Schematic representation of the CRISPR-LICHT screening workflow and experimental design used for the SOX2- and Tdark-focused libraries. Cells were transduced with GFP-expressing lentiviral gRNA libraries before organoid generation and enriched for transduced cells by GFP-based FACS sorting or, where indicated, puromycin selection. Following PSC aggregation, eCas9 activity was induced with 4-hydroxytamoxifen (4-OHT), and organoids were differentiated until day 30–35. At the endpoint, organoids were pooled and dissociated, followed by either dTomato-positive FACS enrichment or direct processing without endpoint enrichment. Genomic DNA was then subjected to barcode-locus amplification, next-generation sequencing and barcode-based lineage analysis. The SOX2-focused library comprised 13 neurodevelopmental regulators targeted by four gRNAs per gene (53 gRNAs in total) and was used to compare two initial enrichment strategies (GFP-based FACS or puromycin selection) and two endpoint processing strategies (dTomato FACS or no endpoint enrichment), generating four experimental conditions: FACS–FACS (I), FACS–NO FACS (II), PURO–FACS (III) and PURO–NO FACS (IV). Twenty organoids were analysed per condition across two replicates, with ten pooled organoids per replicate. The Tdark-focused library targeted 120 candidate SOX2-dependent genes using five gRNAs per gene and included positive and non-targeting control gRNAs, resulting in a total library complexity of approximately 800 gRNAs. The Tdark screen was performed using GFP- based enrichment followed by endpoint dTomato FACS sorting, with 160 organoids distributed across 25 replicates of six to seven pooled organoids each.

The SOX2-focused library was therefore screened under four parallel conditions: initial GFP FACS enrichment followed by “endpoint” (at the end of the organoid culture) dTomato FACS sorting (I, the standard CRISPR-LICHT workflow); initial GFP FACS enrichment followed by bulk endpoint harvest without sorting (II); initial puromycin selection followed by endpoint dTomato FACS sorting (III); initial puromycin selection followed by bulk endpoint harvest These conditions generated four corresponding sequencing datasets: FACS-FACS (I, standard), FACS-NO FACS (II), PURO-FACS (III) and PURO-NO FACS (IV) (Fig. 3B). The SOX2-focused library contained 52 gRNAs targeting *SOX2* and 12 established SOX2- dependent neurodevelopmental regulators, with four gRNAs per gene (Table S3).

In contrast to the Sox2-focused library, the Tdark-focused library was screened using the standard GFP-enrichment and endpoint dTomato-sorting workflow (condition I) and contained approximately 800 gRNAs targeting 120 poorly characterized SOX2-dependent candidate genes, with five gRNAs per gene, together with positive and non-targeting controls (Table S6).

For the SOX2-focused screen, 20 organoids were generated per condition and pooled in groups of 10 organoids, yielding two biological replicates per condition. For the Tdark- focused screen, 160 organoids were generated and pooled in groups of 6–7 organoids, yielding 25 biological replicates (see *Materials and Methods*). Following dissociation, samples were processed according to the corresponding endpoint strategy before genomic DNA extraction, amplification of gRNA-lineage barcode pairs, next-generation sequencing and integrated lineage-resolved analysis (Fig. 3; Table S7).

This experimental design enabled parallel assessment of established SOX2-dependent regulators and poorly characterized candidate genes within a shared organoid-based screening framework, while allowing the SOX2-focused screen to be used to assess the impact of alternative sample-processing strategies on the lineage-resolved readout.

### Evaluation of enrichment and endpoint processing strategies in the SOX2-focused screen

Using the SOX2-focused library, we first examined whether the initial enrichment strategy affected the abundance of “Cas9 activated/expressed” (dTomato reporter-defined) populations after organoid differentiation. Flow cytometry analysis following organoid dissociation before endpoint processing revealed that puromycin selection increased the proportion of dTomato-positive cells detected at the endpoint and reduced the GFP-negative/dTomato-negative fraction compared with standard GFP FACS-based enrichment (Fig. 4A). This suggests that puromycin selection retained a larger fraction of cells carrying transcriptionally active lentiviral integrations, whereas GFP-based sorting may include integrations that subsequently to initial expression undergo silencing during differentiation. The detection of a dTomato-positive/GFP-negative population, consistent with the pilot validation experiment, further supports GFP silencing in a subset of transduced cells rather than inclusion of non-transduced cells (Fig. S3).

**Figure 4.**
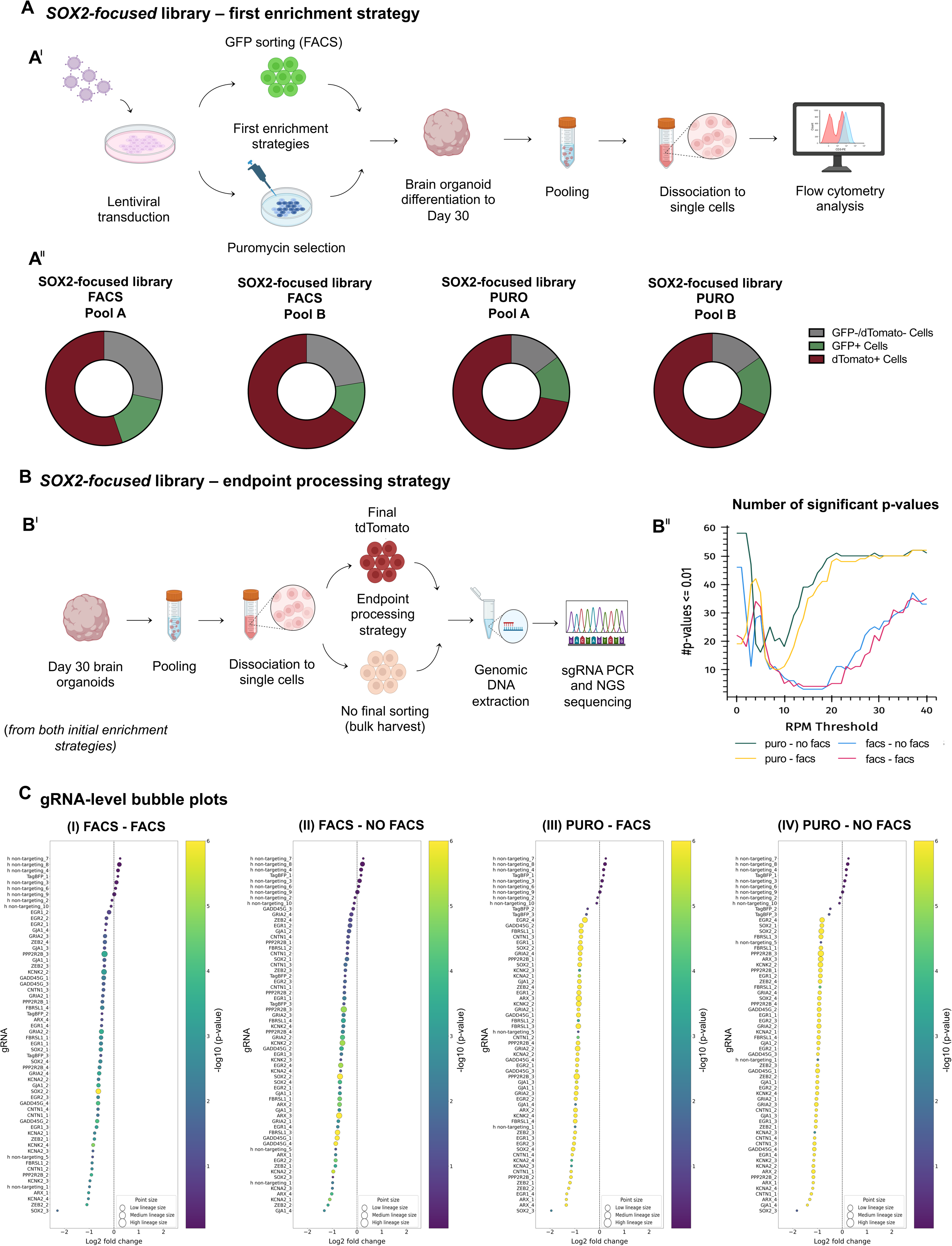
Comparison of initial enrichment and endpoint processing strategies in the SOX2-focused CRISPR screen in day-30 human brain organoids. **(A)** Assessment of initial enrichment strategies for transduced cells. **(I)** Schematic overview of the experimental workflow. hESCs were transduced with the SOX2-focused gRNA library and enriched either by GFP-based FACS sorting or by puromycin selection before organoid generation. Day-30 organoids were pooled, dissociated into single cells and analysed by flow cytometry to quantify reporter-defined populations. **(II)** Relative abundance of GFP-positive, dTomato-positive and GFP-negative/dTomato-negative cells in two independent pools of 10 organoids per condition. dTomato-positive cells indicate activation of inducible eCas9, GFP-positive cells represent transduced cells without detectable eCas9 activation, and GFP- negative/dTomato-negative cells represent cells lacking detectable reporter expression. Compared with GFP-based FACS enrichment, puromycin selection increased the proportion of dTomato-positive cells and reduced the GFP-negative/dTomato-negative fraction, suggesting improved recovery of cells with activated inducible eCas9. **(B)** Comparison of endpoint processing strategies. **(I)** Schematic overview of the two endpoint workflows tested at day 30. Organoids generated after either initial GFP sorting or puromycin selection were pooled and dissociated, followed by either final dTomato-positive FACS sorting or direct bulk harvest without endpoint sorting. Genomic DNA was extracted and gRNA abundance was quantified by PCR amplification and next-generation sequencing. **(II)** Number of significant gRNA-level effects detected across RPM filtering thresholds in the four screening conditions: FACS–FACS, FACS–NO FACS, PURO– FACS and PURO–NO FACS. Significant gRNAs were defined as those showing enrichment or depletion at *P* ≤ 0.01. This analysis was used to assess the impact of read-count filtering and endpoint processing on the detection of significant gRNA-level changes. **(C)** Ranked gRNA-level bubble plots for the four screening conditions: **(I)** FACS–FACS, **(II)** FACS–NO FACS, **(III)**PURO–FACS and **(IV)** PURO–NO FACS. In each plot, gRNAs are ranked by log₂ fold change within the corresponding condition. The dashed vertical line marks log₂ fold change = 0. Dot colour represents −log₁₀(*P* value), capped at 6 for visualization, and dot size indicates lineage size. Non-targeting control gRNAs are included as internal references to visualize background variation and condition-specific shifts in gRNA abundance.

We next asked whether distinct enrichment strategies affected the sequencing-based screen readout in terms of screening sensitivity and recovery of significant gRNA-level effects when combined either with the standard endpoint strategy, consisting of dTomato-positive FACS sorting, or with direct bulk harvesting following organoid dissociation (Fig. 4B). To assess whether the enrichment strategy affected screening sensitivity, defined here as the ability to recover gRNAs targeting known regulators as significant hits, we compared the number of significant gRNAs across conditions under different filtering stringencies. Specifically, we applied increasing reads-per-million (RPM) thresholds to progressively exclude less-represented gRNAs from the analysis. Across RPM thresholds, puromycin-selected conditions yielded a higher number of significant gRNA-level effects than GFP- enriched conditions, particularly at lower filtering thresholds (Fig. 4B). By contrast, within each initial enrichment background, endpoint dTomato FACS sorting had a more limited impact, with sorted and bulk endpoint samples showing broadly comparable numbers of significant gRNAs across thresholds (Table S8). Thus, the initial enrichment strategy had a stronger influence on screening sensitivity than endpoint processing.

To further compare effects at the individual gRNA level, we ranked gRNAs according to their log₂ fold change within each condition (Fig. 4C). Non-targeting gRNAs were retained as internal reference controls to define the background distribution of lineage-size changes across conditions. Across all four conditions, gRNAs targeting SOX2-focused library genes showed predominantly negative log₂ fold changes, indicating reduced lineage size. This pattern was evident for several genes. For example, gRNAs targeting *SOX2* were depleted across all four conditions, consistent with its requirement for neural progenitor maintenance. *ZEB2*, a regulator of early lineage specification, also showed negative fold changes across conditions, but statistical significance was stronger after puromycin selection: two of four *ZEB2*-targeting gRNAs reached *P* ≤ 0.01 in each GFP-enriched condition, whereas all four guides were statistically significant in both puromycin-selected conditions (Fig. 4C; Table S8). A similar pattern was observed for gRNAs targeting *GADD45G* and *KCNA2*, which showed consistent negative effects but more uniform guide-level significance in puromycin-selected samples. Overall, gRNA behaviour was maintained across workflow variants, indicating that the screen captured consistent perturbation-associated depletion phenotypes independently of the specific processing strategy, whereas differences between conditions primarily affected the strength and reproducibility of statistical support.

Together, these data show that the SOX2-focused screen recovered consistent depletion phenotypes for established SOX2-dependent regulators in human brain organoids. Alternative processing strategies did not substantially alter the direction of these effects but modulated the sensitivity of gRNA-level hit detection.

### The Tdark-focused screen identifies poorly characterized regulators of progenitor lineage expansion

Having established the technical performance of the CRISPR-LICHT readout using the SOX2-focused library, we next analyzed the Tdark-focused discovery screen. To better compare the new results with previous work by Esk et al., we used for this analysis the standard FACS-based workflow (Fig. 5A).

**Figure 5.**
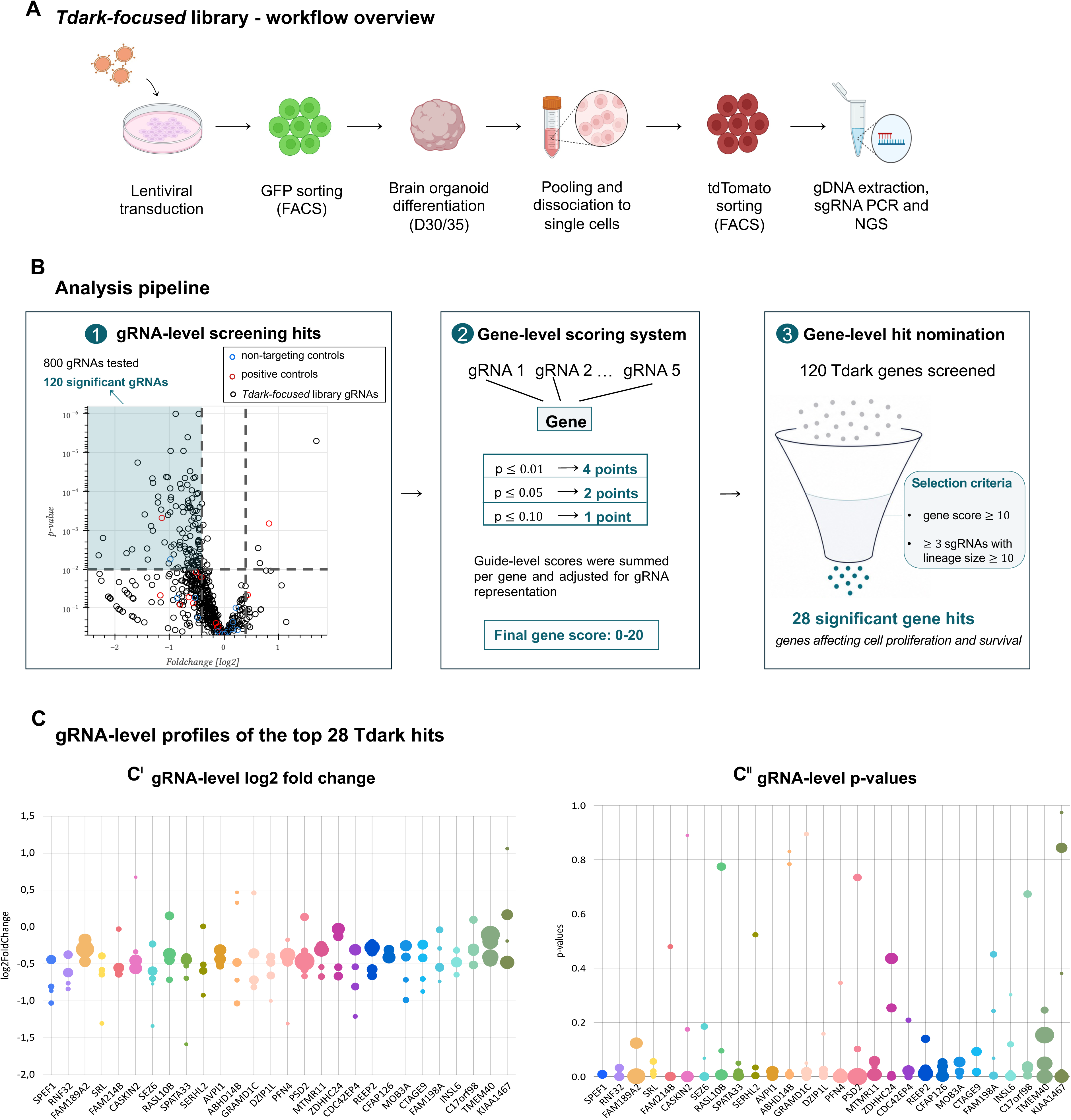
Identification and annotation of candidate SOX2-dependent regulators from the Tdark-focused CRISPR screen. **(A)** Schematic overview of the Tdark-focused CRISPR screening workflow. hESCs were transduced with the Tdark-focused gRNA library, enriched by GFP-based FACS sorting and differentiated into human brain organoids. At day 30/35, organoids were pooled, dissociated into single cells and subjected to dTomato-positive FACS sorting. Genomic DNA was then extracted for gRNA amplification and next-generation sequencing, enabling the identification of candidate genes affecting organoid cell proliferation and/or survival. **(B)** Analysis pipeline used for hit identification from gRNA-level screening data. Individual gRNAs were first evaluated based on their fold change, statistical significance and lineage representation. gRNAs with significant depletion or enrichment and more than 10 detected lineages were retained for downstream analysis. A gene-level score was then calculated by integrating the behaviour of the five independent gRNAs targeting each gene, with each gRNA contributing points according to its statistical significance. This scoring strategy generated a cumulative gene score ranging from 0 to 20 and was used to prioritize genes supported by concordant gRNA-level effects. Genes with a score ≥10 and at least three significant gRNAs with more than 10 lineages were nominated as candidate hits, resulting in 28 Tdark genes predicted to affect organoid cell proliferation and/or survival. **(C)** gRNA-level profiles of the top 28 Tdark candidate hits. **(I)** Distribution of log₂ fold changes for significant gRNAs targeting each of the top 28 genes. Only significant gRNAs represented by more than 10 lineages are shown. Dot size indicates lineage size. **(II)** Distribution of gRNA-level *P* values for the same set of significant gRNAs. In both panels, individual gRNAs are grouped by target gene, allowing visualization of the consistency and strength of the screening phenotype across independent gRNAs. Dot size indicates lineage size.

We retrieved sequences of gRNAs and their relative abundance from endpoint samples, together with plasmid library and timepoint 0 (T0, corresponding to the onset of organoid differentiation) samples. The latter served as technical controls to assess the composition of the library, including the initial gRNA representation in it. These quality-control analyses confirmed maintenance of library complexity without evidence of major bottleneck effects, while also indicating that endpoint lineage abundance was not simply determined by initial gRNA representation (Fig. S5; Fig. S6A; Tables S14, S15).

To focus the analysis on biologically informative lineage-size changes, lineage-level data were filtered using RPM-based thresholds. An RPM threshold of 40 was selected based on the separation of *P*-value distributions between non-targeting control guides and proliferation-associated control guides and was then used for downstream hit identification (Fig. S6). This filtering strategy reduced the contribution of weakly represented or non-proliferative clones while retaining lineages carrying informative proliferation or survival phenotypes (Fig. S6B,C; Table S9).

Using this threshold, we identified 120 gRNAs out of approximately 800 (15%) with significant negative effects on lineage size in endpoint samples, corresponding to perturbations that impaired clonal expansion during organoid differentiation (Fig. 5B). Significant guides showed log₂ fold changes ranging from −2.30 to −0.30, corresponding to lineage sizes of approximately 20–81% of the mean lineage size observed for non-targeting controls. This indicated a broad spectrum of depletion phenotypes, ranging from strong impairment of lineage expansion to more moderate reductions in progenitor fitness (Table S10).

To nominate top hits from the screen, we applied a cumulative scoring strategy integrating statistical significance and concordance across the five gRNAs targeting each gene. This approach prioritized genes showing consistent effects across multiple independent gRNAs rather than candidates driven by a single outlier guide. Briefly, individual gRNAs represented by more than 10 detected lineages were assigned a score according to their significance level, thereby minimizing the contribution of underrepresented perturbations. Guide-level scores were then summed for each gene and adjusted for gRNA representation (see *Materials and Methods*). Candidate hits were selected based on a cumulative gene score ≥10 on a scale from 0 to 20 and with at least three of the five targeting gRNAs meeting the inclusion criterion. Using this scoring framework, we identified 28 hits affecting organoid cell proliferation and/or survival (Fig. 5B). The resulting ranked list of hits is reported in Table S10.

We next visualized the evidence from individual gRNAs underlying the 28 genes nominated by the gene-level scoring approach. Here, “individual gRNA level” refers to the effect measured for each single guide RNA targeting a candidate gene, whereas “gene level” refers to the aggregated score used to nominate genes based on the combined behaviour of multiple gRNAs. For each candidate gene, gRNAs were plotted according to log₂ fold change, *P* value and lineage size (Fig. 5C). This analysis provided a direct view of the individual gRNA effects contributing to gene nomination and showed that, although all 28 genes met the nomination criteria, their profiles differed in the concordance between effect size and statistical support. Some candidates showed consistent negative fold-change trends across multiple gRNAs, whereas others displayed more variable effect sizes. Among the nominated genes, *SPEF1, RNF32, SRL, SEZ6, SPATA33, DZIP1L, CDC42EP4, REEP2, MOB3A* and *CTAGE9* showed particularly concordant gRNA effects, with multiple independent gRNAs displaying negative fold changes together with stronger statistical support (Fig. 5C). These candidates therefore represented the most consistently supported hits at the guide level within the broader gene-level nominated set.

Together, these results identify a set of poorly characterized SOX2-dependent candidate genes required for progenitor lineage expansion and/or survival in human brain organoids.

### Targeted functional follow-up of CDC42EP4 reveals altered neural progenitor distribution in human brain organoids

To prioritize candidates for targeted functional follow-up, we first examined the protein-class composition of the 28 nominated hits. These included signalling proteins, structural or cytoskeletal proteins, scaffold or adaptor proteins, enzymes and membrane-trafficking components, together with poorly characterized or unclassified proteins (Fig. 6A), suggesting potential roles in cellular architecture, cytoskeletal regulation, membrane-associated processes and tissue growth, extending beyond gene regulation.

**Figure 6.**
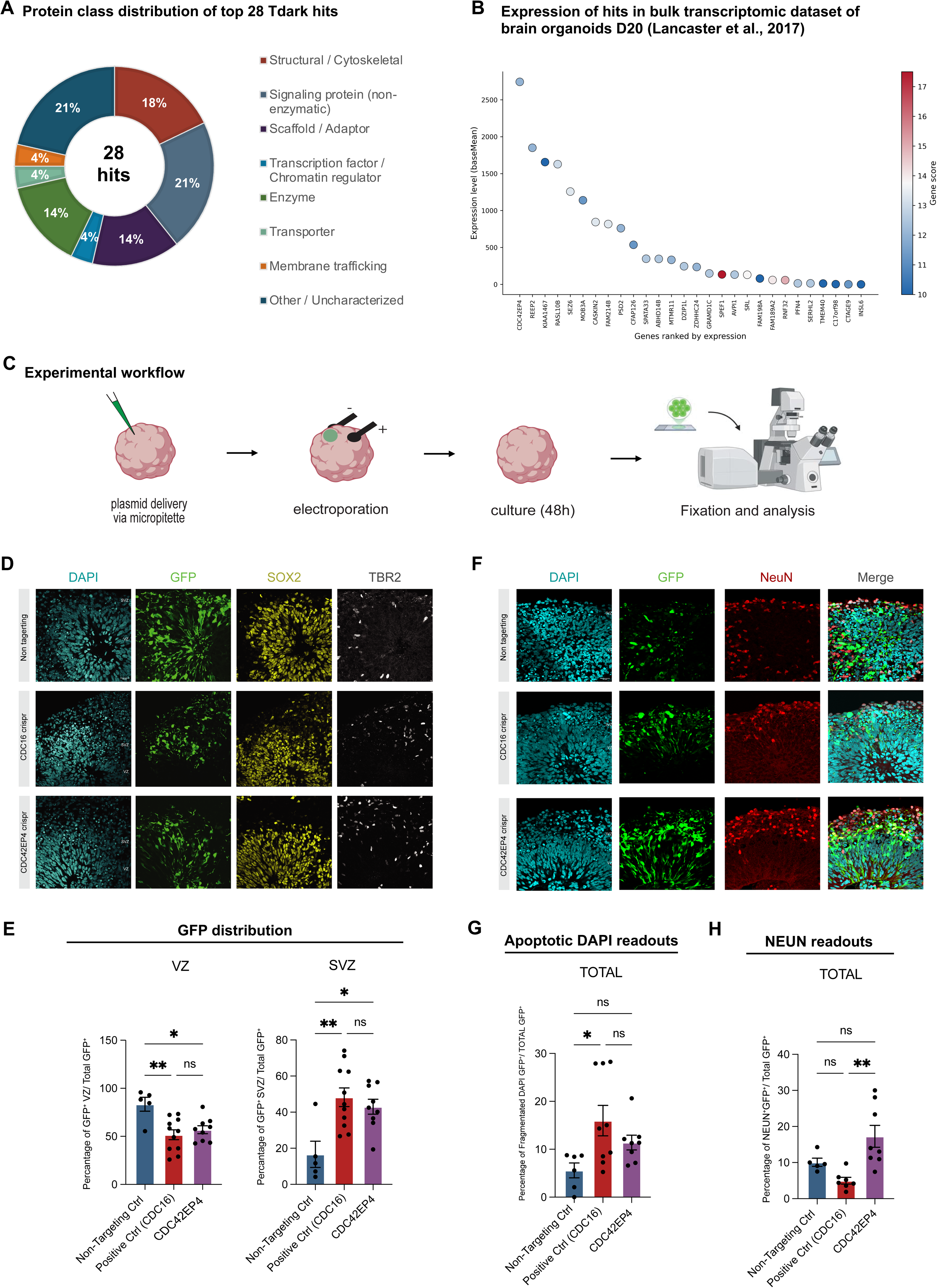
Targeted functional follow-up of CDC42EP4 in human brain organoids. **(A)** Protein-class distribution of the top 28 Tdark candidate hits. Genes were assigned to broad functional categories based on available protein annotation. The highest represented classes included structural/cytoskeletal proteins and signalling proteins, while a substantial fraction of candidates belonged to poorly characterized or unclassified categories, consistent with the Tdark nature of the screened gene set. **(B)** Expression of the 28 top-ranked genes identified in the Tdark-focused CRISPR screen was assessed using the published cerebral organoid dataset from Lancaster *et al*. (2017), generated with a comparable organoid differentiation protocol. Genes are ordered by decreasing expression level, reported as baseMean. Dot color represents the gene score assigned by the screening pipeline, calculated from the aggregate statistical performance of the five gRNAs targeting each gene. All 28 candidates showed a gene score ≥10, while displaying variable expression levels in the organoid dataset. Integration of screen-derived gene scores with published organoid expression data was used to prioritize candidates for downstream functional follow-up. **(C)** Schematic representation of the electroporation-based follow-up experiment. H9 cortical organoids were electroporated at D35 with a non-targeting control guide, the CDC16 positive-control CRISPR guide, or the CDC42EP4 CRISPR guide, and analyzed at D37. **(D)** Representative immunofluorescence images of electroporated ventricular regions from H9 organoids, electroporated at D30 and analyzed at D32 with control (non- targeting guide), CDC16 CRISPR guide, and CDC42EP4 CRISPR guide cortical organoids. Sections were stained for DAPI (cyan), GFP (green), SOX2 (yellow), and TBR2 (white). Ventricular zones (VZ) and subventricular zones (SVZ) were identified based on DAPI staining and tissue organization. Scale bar, 20 μm. **(E)** Quantification of GFP^+^ cells within the VZ and SVZ. Graphs show the percentage of GFP-positive cells in the indicated regions, relative to the total number of GFP- positive cells. Data are presented as mean ± SEM. Each dot represents an individual ventricle analyzed. A total of 5–11 electroporated ventricles from 6-7 independent organoids from two independent batches were analyzed per condition. Statistical significance was determined using one-way ANOVA with Turkey’s multiple comparisons test. ns, not significant; *P < 0.05; **P < 0.01. **(F)** Representative immunofluorescence images of electroporated ventricular regions fromH9 organoids, electroporated at D30 and analyzed at D32 with control (non- targeting guide), CDC16 CRISPR guide, and CDC42EP4 CRISPR guide cortical organoids. Sections were stained for DAPI (cyan), GFP (green), NeuN (red), and a merged image of all the channels. Ventricular zones (VZ) and subventricular zones (SVZ) were identified based on DAPI staining and tissue organization. Scale bar, 20 μm. **(G)** (Left) Quantification of the percentage of NEUN-positive cells among GFP-positive cells within ventricular regions. Data are shown as mean ± SEM. Each dot represents an individual ventricle analyzed. A total of 5-8 electroporated ventricles from 6-7 independent organoids from two independent batches were analyzed per condition. Statistical significance was determined using one-way ANOVA with Turkey’s multiple comparisons test, ns, not significant; **P < 0.01. (Right) Quantification of cells fragmentated DAPI and are GFP^+^ along the electroporated apical surface of cortical organoid ventricles. The number of apoptotic cells was normalized to 100 μm of electroporated apical surface. Bars represent mean ± SEM, and individual dots represent independent electroporated ventricles. n = 6–9 electroporated ventricles from 6–7 independent organoids from two independent batches were analyzed per condition. Statistical significance was determined using one-way ANOVA followed by Turkey’s multiple comparisons test. ns, not significant; *P < 0.05.

We next assessed the expression of the 28 hits using transcriptomic data from a published cerebral organoid gene expression dataset from organoids generated with a comparable differentiation protocol and analyzed at a similar developmental stage (Div 20; Lancaster et al., 2017) (Fig. 6B), together with data from the developing human brain from the Allen Human Brain Atlas (Fig. S7; Table S17). All 28 candidates were expressed, although at variable levels across the two datasets. Notably, *CDC42EP4* was the most highly expressed hit in the cerebral organoid dataset and showed consistent expression in the developing human brain (as represented in the Allen Human Brain Atlas), supporting its prioritization for functional investigation. Available annotations link CDC42EP4 to CDC42-related cytoskeletal regulation, including actin-associated processes, cell-shape control and cellular polarity. We therefore selected *CDC42EP4* for targeted functional follow-up in human cerebral organoids.

To assess the consequences of *CDC42EP4* perturbation on neural progenitor abundance and distribution, we established an independent CRISPR-based electroporation assay in cerebral organoids (Fig. 6C). Cas9 together with sgRNAs targeting *CDC42EP4* were electroporated into ventricular zone-like (VZ-like) structures of div 35 organoids generated from the H9 hESC line, thereby targeting spatially defined progenitor domains. Organoids were electroporated with either a non-targeting sgRNA, a *CDC16*-targeting sgRNA used as a positive control, or a *CDC42EP4*-targeting sgRNA and analyzed 48 h after electroporation. *CDC16*, a core component of the anaphase-promoting complex/cyclosome (APC/C) required for cell-cycle progression (King et al., 1995), was included as a reference perturbation expected to impair progenitor expansion. To assess the spatial distribution and cellular identity of electroporated cells, we performed immunostaining for GFP together with SOX2 and TBR2 in cryosections of electroporated organoids (Fig. 6D). TBR2 marks intermediate neural progenitors associated with progression from ventricular radial glial progenitors toward a neurogenic lineage (Englund et al., 2005; Hevner, 2019) and was therefore included to assess whether changes in spatial distribution were accompanied by altered progenitor-state composition. GFP-positive cells were quantified within VZ-like regions and adjacent subventricular zone-like (SVZ-like) regions. In the non-targeting condition, GFP-positive cells were predominantly located within the VZ-like compartment, in agreement with the position the DNA delivery. *CDC16* perturbation reduced the proportion of GFP-positive cells within the VZ-like region and increased their representation in the SVZ-like compartment, possibly suggesting that their survival or proliferation had been decreased.

*CDC42EP4* perturbation produced a similar spatial redistribution: *CDC42EP4*-targeted GFP-positive cells were significantly less represented within the VZ-like compartment and significantly more represented within the SVZ-like compartment compared with the non-targeting condition (Fig. 6E). Thus, perturbation of *CDC42EP4* in this orthogonal assay altered the distribution of targeted cells between ventricular- and subventricular-like compartments.

We next asked whether this spatial redistribution was accompanied by loss of SOX2-positive progenitor identity. GFP-positive cells remained predominantly SOX2-positive following *CDC42EP4* perturbation, and quantitative analysis did not reveal a significant change in the overall proportion of SOX2-positive cells among GFP-positive cells compared with either the non-targeting or *CDC16*-targeting conditions (Fig. S8). Thus, redistribution of *CDC42EP4*-targeted cells occurred while SOX2 expression was largely maintained over the 48-h experimental window, consistent with substantial retention of progenitor identity at this time point.

The redistribution of targeted cells from VZ-like to SVZ-like regions could reflect reduced survival within the VZ-like compartment or progression toward more differentiated neurogenic states. To distinguish between these possibilities, we analyzed apoptosis together with the intermediate progenitor marker TBR2 and the neuronal marker NeuN (Fig. 6F,G,H). *CDC16* perturbation resulted in a significant increase in apoptotic cells (pyknotic nuclei by DAPI staining) compared with the non-targeting condition, consistent with reduced survival following cell-cycle disruption (Fig. 6G; Fig. S8). In contrast, *CDC42EP4* perturbation did not result in a statistically significant increase in apoptosis.

TBR2-positive cells showed some (not statistically significant) increase following *CDC42EP4* perturbation, particularly within the SVZ-like compartment (Fig. S8). Similarly, the proportion of NeuN-positive cells among GFP-positive cells was somewhat increased, though not in a significantly statistical way, following *CDC42EP4* perturbation compared with the non-targeting condition (Fig. 6H). NeuN positivity was, however, significantly higher following *CDC42EP4* perturbation than following *CDC16* perturbation (Fig. 6H). Together, these findings indicate that, despite producing a spatial redistribution similar to that observed following *CDC16* perturbation, *CDC42EP4* targeting was not associated with increased apoptosis, and instead showed trends toward initial increased differentiation, as indicated by the larger proportion of both intermediate progenitor and neuron markers. These results are consistent with altered progenitor-state progression following *CDC42EP4* perturbation rather than depletion driven primarily by cell death.

## Discussion

In this study, we applied CRISPR-LICHT screening in human cerebral organoids to functionally interrogate 133 SOX2-associated genes during early neural development. By combining a SOX2-focused reference library with a discovery screen targeting poorly characterized Tdark genes, we identified novel regulators of human NSC abundance and uncovered a broader spectrum of cellular processes potentially associated with SOX2- dependent neural progenitor biology. In particular, the Tdark screen highlighted genes linked to signalling, cytoskeletal organization and membrane-associated processes, suggesting that SOX2-associated developmental programmes may extend beyond transcriptional control of neural identity to encompass cellular mechanisms supporting progenitor organization and expansion. Targeted functional analysis of CDC42EP4 further supports this view, identifying a poorly characterized CDC42 effector whose perturbation alters the spatial distribution and developmental state of neural progenitor populations in human cerebral organoids.

The SOX2-focused screen provided a reference for assessing CRISPR-LICHT in the context of SOX2-associated neural progenitor regulation. SOX2 perturbation strongly reduced lineage contribution across the experimental conditions tested, consistent with its central role in progenitor maintenance. Depletion phenotypes were also observed for genes with diverse established neurodevelopmental functions, including regulators of early patterning and fate specification such as ZEB2 and ARX (Garavelli et al., 2017; Nieto-Estevez et al., 2024), the proliferation- and differentiation-associated factor GADD45G (Patel et al., 2022), and genes more commonly linked to neuronal maturation and excitability, including KCNA2 and GRIA2 (Syrbe et al., 2015; Efthymiou et al., 2024). Although these candidates were selected because of their association with SOX2-dependent transcriptional programmes and neurodevelopmental function, their convergent effects on lineage contribution during an early progenitor-enriched developmental window are notable. In particular, depletion of genes best characterized for later neuronal functions, such as GRIA2, suggests that their contribution to neural development may begin earlier than indicated by their canonical annotations. Such early requirements could reflect additional functions in progenitor-intrinsic processes that are later reutilized during neuronal maturation, or early perturbations whose consequences become progressively amplified during lineage expansion. Because lineage size integrates the cumulative consequences of gene perturbation on proliferation, survival and developmental progression, the present screen cannot resolve the cellular mechanism underlying each phenotype. Nevertheless, these results illustrate how lineage-resolved functional screening can reveal developmental requirements that are not readily predicted from the best-characterized functions of individual genes.

The SOX2-focused library also allowed us to assess how experimental processing influences the sensitivity of lineage-resolved CRISPR-LICHT screening. Although the overall direction of guide-level effects was broadly maintained across workflows, supporting the methodological robustness of the screening platform, puromycin-based enrichment increased the recovery and statistical support of depletion phenotypes compared with GFP- based sorting, likely reflecting, at least in part, improved lineage representation following selection. By contrast, endpoint dTomato sorting had a comparatively limited influence on the major phenotypic patterns. Importantly, robust lineage-depletion phenotypes could also be recovered from bulk endpoint samples without final dTomato enrichment, indicating that, at least for sufficiently strong phenotypes, CRISPR-LICHT screening can be performed without endpoint cell sorting. Eliminating this step could simplify the experimental workflow, reduce cell loss during sample processing and facilitate applications in laboratory settings in which sorting capacity or experimental throughput is limiting. Together, these observations suggest that optimizing the initial enrichment of transduced cells may have a greater impact on screening sensitivity than endpoint purification, providing practical guidance for future applications of CRISPR-LICHT.

Building on this reference screen, the Tdark-focused library was designed to move beyond well-characterized neurodevelopmental regulators and interrogate genes with limited functional annotation. The identification of 28 nominated candidates indicates that this poorly explored gene space contains regulators whose perturbation measurably affects lineage contribution during early human brain organoid development. Importantly, although the targeted genes encompassed heterogeneous protein classes, several nominated candidates were associated with signalling, cytoskeletal organization, membrane dynamics and cellular architecture. These functions are particularly relevant in neural progenitors, whose behaviour depends on coordinated apical-basal polarity, ventricular organization, cell-cycle progression and the balance between progenitor maintenance and differentiation (Andrews et al., 2022).

This convergence on structural and spatial processes may represent a less explored component of SOX2-associated biology. While substantial progress has been made in characterizing SOX2-dependent transcriptional networks involved in maintaining neural progenitor identity (Pevny and Nicolis, 2010; Shimozaki, 2014), comparatively less is known about how these programmes connect to cellular architecture. Recent work in the mouse pre-implantation epiblast identified enrichment of actin cytoskeletal, cell-cell junction, focal adhesion and extracellular-matrix functions among genes associated with putative SOX2 binding and showed that Sox2 loss alters epithelial polarization and tissue organization (Chandramohan et al., 2026). Although this represents a distinct developmental context, it supports the broader possibility that SOX2 may coordinate transcriptional cell state with mechanisms controlling cellular architecture. The Tdark candidates identified here suggest that similar relationships may operate within neural progenitor populations, where structural and membrane-associated effectors could contribute to the organization and proliferative capacity of ventricular progenitor compartments.

Among the nominated candidates, CDC42EP4 emerged as a compelling candidate for functional follow-up because of its strong expression in cerebral organoids and the developing human brain, together with its poorly characterized role in neurodevelopment. Our electroporation experiments showed that targeted perturbation of CDC42EP4 within VZ-like regions produced a clear redistribution of electroporated cells, with reduced representation within the VZ-like compartment and increased representation within the adjacent SVZ-like region. Importantly, CDC42EP4-targeted cells largely retained SOX2 expression, indicating that this spatial phenotype occurred within a population that maintained a major neural progenitor marker over the experimental window. Together, these observations raise the possibility that CDC42EP4 contributes to coupling progenitor state with spatial organization within VZ-like structures.

The comparison with CDC16 further suggests that superficially similar changes in progenitor distribution may arise through different cellular processes. Both CDC16 and CDC42EP4 perturbation reduced the representation of targeted cells within VZ-like regions and increased their representation in SVZ-like compartments. However, CDC16 targeting was associated with a significant increase in apoptotic cells relative to the non-targeting condition, consistent with the consequences expected from disruption of a regulator of cell-cycle progression. In contrast, CDC42EP4 perturbation did not produce a statistically significant increase in apoptosis. TBR2-positive cells also showed a modest, non-significant increase following CDC42EP4 perturbation, particularly within the SVZ-like compartment, consistent with a possible shift toward a more neurogenic progenitor state. More notably, NeuN-positive cells were more abundant following CDC42EP4 perturbation than following CDC16 targeting and also showed a numerical increase relative to the non-targeting condition, although the latter comparison did not reach statistical significance. Although these data do not establish increased neuronal differentiation as the mechanism underlying the CDC42EP4 phenotype, their combined pattern raises the possibility that altered progenitor progression, rather than increased cell death alone, contributes to the reduced representation of CDC42EP4-targeted cells within the VZ compartment.

One possible mechanism linking CDC42EP4 to this phenotype involves cytoskeletal regulation via CDC42. CDC42EP4 belongs to a family of CDC42 effectors associated with septin- and cytoskeleton-related processes (Ageta-Ishihara et al., 2018), whereas CDC42 itself has established roles in epithelial and neuroepithelial polarity, radial glial organization and early brain morphogenesis (Andrews et al., 2022). Perturbation of CDC42-associated cytoskeletal regulation could therefore influence the positioning, maintenance or developmental progression of neural progenitors within ventricular structures, a possibility that is particularly intriguing in light of the spatial redistribution observed following CDC42EP4 targeting. These observations support a working model in which CDC42EP4- dependent cytoskeletal regulation contributes to the organization and developmental progression of neural progenitors within ventricular compartments (Fig. 7). Interestingly, CDC42EP4 plays important roles in Bergmann glia of the mouse cerebellum, where also SOX2 plays an important function (Cerrato et al., 2018). However, direct evidence for CDC42 involvement, altered CDC42 activity, changes in cytoskeletal organization or disruption of apical-basal polarity is not provided by the present study. Rather, the progenitor phenotype identified here establishes a neurodevelopmental context in which the molecular functions of CDC42EP4 can now be investigated. Further analysis will be required to define the mechanisms through which CDC42EP4 contributes to progenitor behaviour.

**Figure 7.**
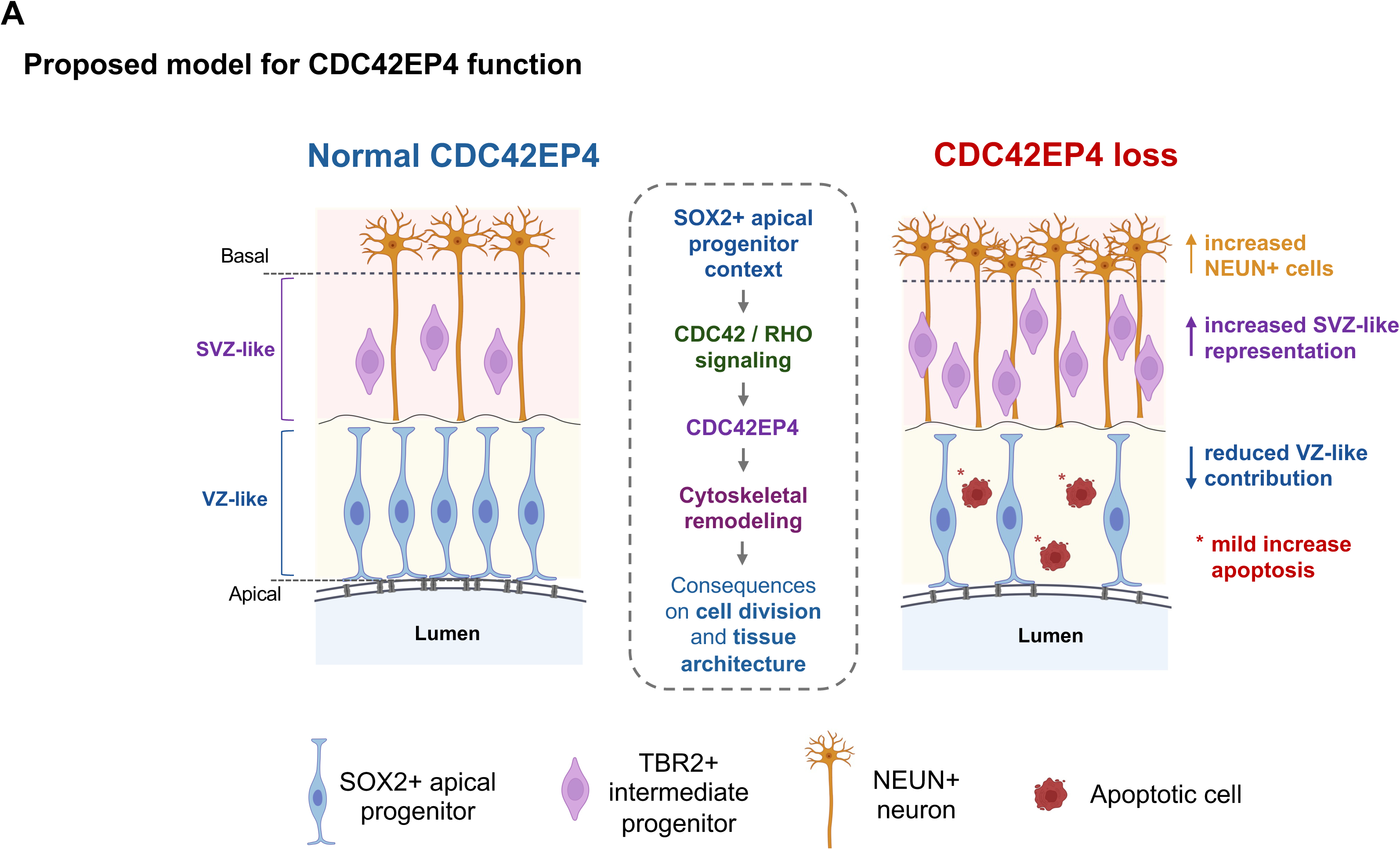
Proposed model for CDC42EP4 function in human apical progenitors. **(A)** Proposed model for CDC42EP4 function in ventricle-like regions of human brain organoids. In the presence of normal CDC42EP4 function, SOX2- positive apical progenitors are maintained within the VZ-like compartment adjacent to the lumen, while TBR2-positive intermediate progenitors and NEUN-positive neurons occupy progressively more basal regions. Following CDC42EP4 loss, the model depicts reduced VZ-like progenitor contribution, increased SVZ-like representation, increased NEUN-positive neuronal differentiation and a mild increase in apoptosis. The central putative pathway places CDC42EP4 downstream of CDC42/RHO signaling in a SOX2-positive apical progenitor context, where it may contribute to septin/cytoskeletal organization and thereby influence cell shape, adhesion and progenitor-state progression. This model is consistent with the reduced lineage contribution observed upon CDC42EP4 targeting and with the increased neuronal differentiation observed in candidate-focused perturbation experiments.

More broadly, the identification of CDC42EP4 within a SOX2-informed screening framework provides a potential entry point into the relationship between transcriptional progenitor programmes and cellular architecture. Evidence from other systems supports links between SOX2-dependent states and cytoskeletal or adhesive processes. In the early mouse embryo, Sox2-associated genes have been linked to focal adhesion, cell-cell junction and actin cytoskeletal functions, together with regulation of epithelial morphogenesis (Chandramohan et al., 2026), while SOX2 depletion in glioma cells has been associated with altered adhesion, actin organization and RhoA-ROCK signalling (Oppel et al., 2011). Although these systems differ substantially from developing neural progenitors, they are consistent with the possibility that SOX2-associated developmental programmes interface with pathways controlling cellular structure and tissue organization. Within this framework, CDC42EP4 represents a candidate effector through which cytoskeletal regulation may influence neural progenitor behaviour.

### Limitations of the study

Several limitations should be considered when interpreting these findings. First, CRISPR- LICHT measures lineage contribution but does not by itself distinguish the cellular mechanisms responsible for lineage depletion. In addition, CRISPR-based perturbation does not provide direct control over the nature or efficiency of mutations generated at each target locus, and incomplete or heterogeneous loss of function may attenuate some phenotypes. CRISPR-LICHT should therefore be viewed primarily as a functional discovery strategy for nominating regulators of progenitor behaviour, with targeted experiments required to resolve the mechanisms underlying individual hits.

These considerations also apply to CDC42EP4. Electroporation enabled spatially resolved analysis of CDC42EP4-targeted cells within ventricular-like structures, but represents a mosaic perturbation and captures the phenotype over a relatively restricted developmental window. Stable or inducible loss-of-function approaches, together with rescue experiments, will be important to establish phenotype specificity and determine how CDC42EP4 influences progenitor organization and developmental progression. Likewise, direct analysis of CDC42 activity, cytoskeletal dynamics and apical-basal polarity will be required to test the mechanistic model suggested here. The present data also provide suggestive rather than definitive evidence for altered neuronal differentiation following CDC42EP4 perturbation; longer-term lineage analysis will be required to determine whether the redistribution of targeted cells is accompanied by premature or increased neuronal differentiation.

A further consideration is that the candidate libraries were informed by genes whose expression was previously found to depend on Sox2 in murine neural stem cells. Although this strategy enabled functional testing of these candidates in a human organoid context, the present study does not establish that all nominated genes are directly regulated by SOX2, or that their SOX2 dependence is conserved in human neural progenitors. The findings should therefore be interpreted as defining functional candidates emerging from a SOX2-informed framework rather than a direct human SOX2 regulatory network.

Finally, all experiments were performed within the H9 genetic background and focused on relatively early stages of organoid development. The pooled CRISPR-LICHT experiments used H9-derived Cas9-expressing cells, whereas the candidate-focused electroporation experiments were performed using parental H9 cells. Although this provides a controlled genetic context for integrating pooled screening with spatially resolved functional analysis, it does not establish whether the identified phenotypes are conserved across human genetic backgrounds. Similarly, the pooled screen captured cumulative effects on lineage contribution during early organoid development, whereas the CDC42EP4 electroporation assay provided a short-term spatial and cellular snapshot. Testing additional pluripotent stem cell backgrounds and developmental windows will therefore be important for determining the generalizability and stage specificity of the identified gene functions.

## Supporting information

Pozzolini et al Figures S1-S8

Pozzolini et al Tables S1-S17

## Acknowledgements

We thank Jürgen Knoblich (Institute of Molecular Biotechnology of the Austrian Academy of Sciences (IMBA), Vienna, Austria) for providing the eCas9-dTomato knock-in cell line; Marta Suarez Cubero (Institute of Molecular Biology & CMBI, Department of Genomics, Stem Cell & Regenerative Medicine, University of Innsbruck, Innsbruck, Austria), Francesca Mingozzi and Stefania Citterio (Department of Biotechnology and Biosciences, University of Milano-Bicocca, Milan, Italy) for assistance with FACS analysis; Luca Muzio (Institute of Experimental Neurology, Division of Neuroscience, IRCCS Ospedale San Raffaele, Milan, Italy) for methodological support and for sharing expertise and protocols for lentiviral production and use; the facilities of Human Technopole (Milan) for DNA sequencing and genome integrity assays: Nicolò Alfano and Clelia Peano (Human Technopole, Milan, Italy) for high-throughput sequencing of the libraries generated for the CRISPR-LICHT screens and for technical advice and Gedi facility (Human Technopole, Milan, Italy) for genome integrity analysis; Sara Barilà (Department of Biotechnology and Biosciences, University of Milano-Bicocca, Milan, Italy) for helping with cerebral organoid cryosectioning, IF and imaging. We also thank the laboratory of Sara Mercurio (Department of Biotechnology and Biosciences, University of Milano-Bicocca, Milan, Italy), the laboratories of the Brain4sight EU consortium, and Sergio Ottolenghi, together with our own laboratories, for helpful discussions.

## Materials and Methods

### Cell culture and inducible eCas9-dTomato hESC line

Wild-type H9 human embryonic stem cells (hESCs) were used for parallel experiments and electroporation-based validation assays (WiCell, Madison, WI, USA). H9 hESCs carrying an inducible eCas9-dTomato cassette were obtained from Jürgen Knoblich (IMBA) (Esk et al, Science, 2020), following MTA procedures. The inducible eCas9-dTomato line was used for CRISPR-LICHT screening experiments. In this line, enhanced Cas9 (eCas9) expression is controlled by a Cre-ERT2/lox-STOP-lox system and is coupled to a dTomato fluorescent reporter, enabling identification of cells in which eCas9 expression has been induced following 4-hydroxytamoxifen (4-OHT) treatment.

Cells were maintained on hESC-qualified Matrigel (Corning, 354277) in Essential 8 medium (Thermo Fisher Scientific, A1517001) at 37 °C in 5% CO₂. Cultures were routinely tested for mycoplasma contamination once per month using the N-GARDE Mycoplasma PCR Reagent Set (Euroclone, EMK090020), according to the manufacturer’s instructions, and were confirmed to be negative. Genome integrity of both hESC lines was assessed using a copy-number assay. No copy-number variants were detected in wild-type H9 hESCs, whereas three variants were identified in the inducible eCas9-dTomato line, as reported in Table S16. These variants were not predicted to affect the screening outcomes.

### Brain organoid generation for CRISR-Licht

Brain organoids were generated using a modified version of the Lancaster et al. (2013) protocol (Fig. S1). Briefly, hESCs were grown to 60–80% confluence, dissociated into single cells using Accutase (Sigma-Aldrich, A6964) for 5 min at 37 °C, pelleted and resuspended in Essential 8 medium supplemented with 50 µM ROCK inhibitor Y27632 (Tocris, 1254). Embryoid bodies were initiated by seeding 10,000 cells per well into 96-well ultra-low-attachment U-bottom plates (Corning, 3307) in 150 µL of Essential 8 medium supplemented with ROCK inhibitor (50 µM).

On day 3, medium was replaced with Essential 8. From day 6 to day 10, embryoid bodies were cultured in neural induction medium, consisting of DMEM/F12 (Invitrogen, 11330-032) supplemented with N2 (100x, Invitrogen, 17502048), GlutaMAX (100x, Invitrogen, 35050- 038), MEM non-essential amino acids (100x, Sigma-Aldrich, M7145) and heparin (1mg/ml, Sigma-Aldrich, H3149). On day 10, batches in which more than 80% of embryoid bodies displayed round morphology, smooth edges and peripheral tissue clearing were transferred to 10 cm dishes rinsed with Anti-adherence solution (STEMCELL Technologies, 07010) in neural induction medium supplemented with penicillin/streptomycin (100x, Sigma-Aldrich, P0781) and 2% v/v growth factor-reduced Matrigel (Corning, 356230).

From day 13, organoids were cultured in improved differentiation medium without vitamin A (iDIFF-A), consisting of DMEM/F12 and Neurobasal medium (Invitrogen, 21103049) supplemented with N2 (200x, Invitrogen, 17502048), B27 without vitamin A (50x, Invitrogen, 12587010), human insulin (10mg/ml, Sigma-Aldrich, 113776497001), 2-mercaptoethanol (143 mM), GlutaMAX (100x, Invitrogen, 35050-038), MEM non-essential amino acids (200x, Sigma-Aldrich, M7145) and penicillin/streptomycin (100x, Sigma-Aldrich, P0781). Organoids were exposed to 3 µM CHIR99021 (Bio-Techne, 4423) on days 13 and 14, maintained in iDIFF-A and transferred to orbital shaking culture (62 routes per minutes) from day 19. From day 25, organoids were maintained in complete improved differentiation medium (ciDIFF), consisting of DMEM/F12 and Neurobasal medium supplemented with N2 (100x, Invitrogen, 17502048), B27 with vitamin A (50x, Invitrogen, 17504044), human insulin (10mg/ml, Sigma-Aldrich, 113776497001), 2-mercaptoethanol (143 mM), GlutaMAX (100x, Invitrogen, 35050-038), MEM non-essential amino acids (200x, Sigma-Aldrich, M7145), sodium bicarbonate (Sigma-Aldrich, S5761), vitamin C (0,4 mM, Sigma-Aldrich, A4544) and penicillin/streptomycin (100x, Sigma-Aldrich, P0781), with medium changes every 3–4 days. For experiments requiring eCas9 induction, cultures were treated on day 5 with 0.3 µg/mL 4-OHT (Sigma-Aldrich, H7904) in Essential 8 medium overnight, followed by return to standard differentiation conditions.

Organoid differentiation was monitored by morphology, including the appearance of neuroepithelial structures and rosette-like regions, and by immunostaining of sectioned organoids. For immunofluorescence, organoids were fixed in 4% paraformaldehyde in PBS for 30 min at room temperature, washed three times in PBS and incubated overnight at 4 °C in 30% sucrose in PBS. Tissues were embedded in 7.5% gelatin/10% sucrose in PBS, mounted in OCT compound (Sakura, 4583), frozen in 2-methylbutane (Sigma-Aldrich, M32631) and stored at −80 °C until sectioning. Frozen organoid blocks were cryosectioned at 20 µm and collected on Superfrost Plus slides (Catalog. No. 6776214). Sections were rinsed in PBS, blocked for 1h at room temperature in blocking solution containing 10% normal donkey serum and 0.5% Triton X-100, and incubated with primary antibodies diluted in antibody solution containing 10% normal donkey serum and 0.1% Triton X-100. Primary antibodies used for organoid characterization were SOX2 (mouse, R&D Systems, MAB2018R, 1:200), PAX6 (sheep, R&D Systems, AF8150, 1:200) and FOXG1 (rabbit, Abcam, ab18259, 1:300).

After washing, sections were incubated for 1h at room temperature with donkey secondary antibodies diluted in antibody solution: anti-rabbit Alexa Fluor 647 (Invitrogen, A32795TR, 1:1000), anti-sheep Alexa Fluor 568 (Invitrogen, A21099, 1:1000) and anti-mouse Alexa Fluor 647 (Invitrogen, A32787TR, 1:1000). Nuclei were counterstained with DAPI (MERCK, D9542, 2 µg/mL), and sections were mounted with DAKO Fluorescent Mounting Medium (Dako, S3023). Images were acquired using a THUNDER Imager Tissue Live microscope (Leica) and LASX software using 10x and 20x objectives. Images compared within the same experiment were acquired using identical settings and processed uniformly. Differentiation quality and regional identity were assessed by expression of neural progenitor and dorsal forebrain-associated markers, including SOX2, PAX6 and FOXG1, as shown in Fig. 2 and Fig. S1.

### Brain organoid generation for electroporation experiments

For electroporation experiments, organoids were generated using a modified minimally guided brain organoid differentiation protocol based on Lancaster *et al*. (2014) and Lancaster and Knoblich (2014). Briefly, 9,000 H9 ESCs (WiCell, Madison, WI, USA) were seeded per well in round-bottom 96-well plates (Corning, CLS7007) to induce embryoid body formation and early germ-layer differentiation. From day 5 to day 7, embryoid bodies were cultured in neural induction medium to promote neural ectoderm formation. On day 7, organoids were embedded in Matrigel droplets (Corning, 354234) and transferred to expansion medium to support the formation and expansion of neuroepithelial buds. From day 7 onward, Matrigel-embedded organoids were maintained on an orbital shaker. Organoids were cultured in neural differentiation medium supplemented with B27 without vitamin A (100x, Thermo Fisher Scientific, 12587010) until day 14, and then switched to neural differentiation medium containing B27 with vitamin A (100x, Thermo Fisher Scientific, 17504044). Organoids were maintained in 10-cm dishes on an orbital shaker at 37 °C, 5% CO₂ and ambient oxygen, with medium changes twice per week.

### CRISPR-LICHT library design and cloning Lineage barcode library

A random 10N lineage barcode library was synthesized with flanking sequences compatible with the NheI-HF (New England Biolabs, R3131M) and EcoRI-HF (New England Biolabs, R3101M) cloning sites of the human screening CRISPR-LICHT vector (hSCR), as described by Esk et al. The barcode oligonucleotide library was amplified by qPCR and ligated into NheI/EcoRI-digested hSCR vector using T4 DNA ligase (New England Biolabs, M0202L), generating a pooled lineage barcode library for clonal tracking during organoid development (Fig. S2).

### SOX2-focused and Tdark-focused gRNA libraries

Two CRISPR-LICHT gRNA libraries were designed for this study. The SOX2-focused library targeted SOX2 together with 12 additional SOX2-regulated neurodevelopmental genes previously identified in murine neural stem cells and selected based on established or predicted roles in neural progenitor regulation and early neurodevelopment. Four gRNAs were included per gene, resulting in 52 gRNAs in total.

The Tdark-focused library targeted 120 poorly characterized SOX2-dependent candidate genes selected among evolutionarily conserved transcripts downregulated following *Sox2* loss in murine neural stem cells and classified as Tdark according to the Illuminating the Druggable Genome framework. Five gRNAs were included per gene. The library also contained positive-control and non-targeting control gRNAs, resulting in an overall complexity of approximately 800 gRNAs. All gRNA sequences included in the library were selected from the gRNA collection described by Michlits et al. (2020). Complete library compositions are reported in Tables S3 and S6.

Guides were synthesized with a G(N₁₉) structure and cloned into BbsI-HF-digested hSCR vector (New England Biolabs, R3539M) by Golden Gate assembly. Ligation products were transformed into MegaX DH10 T1R Electrocomp cells (Thermo Fisher Scientific, C640003) according to the manufacturer’s instructions and plated onto LB agar plates containing ampicillin. Dilution series were performed to estimate library complexity, and each pooled library was confirmed to contain at least 1 × 10⁶ colonies. Colonies were scraped from overnight plates and recovered for 1 h in LB medium containing ampicillin. Plasmid DNA was extracted using the EndoFree Plasmid Maxi Kit (Macherey-Nagel, 740424.50).

### Lentiviral production and hESC transduction Lentiviral library production

Lentiviral particles were produced in HEK293T cells maintained in DMEM supplemented with 10% fetal bovine serum, 1% L-glutamine, 100 U/mL penicillin and 0.1 mg/mL streptomycin. Cells were co-transfected with the pooled CRISPR-LICHT plasmid library, psPAX2 packaging plasmid (Addgene, 12260) and VSV-G envelope plasmid (Addgene, 14888) using a standard calcium phosphate transfection protocol. Medium was replaced with fresh growth medium 12 h after transfection. Viral supernatants were collected 60 h after transfection, filtered through 0.45 µm PVDF filters and concentrated by centrifugation at 11,000 rpm for 8h at 4 °C. Viral pellets were resuspended in PBS, aliquoted and stored at −80 °C. Viral titre was determined before hESC transduction to ensure infection at the desired multiplicity of infection. It was estimated by transducing HEK cells with serial dilutions of viral supernatant and quantifying GFP-positive cells by flow cytometry 48h after infection.

### hESC transduction

hESCs were transduced in stem cell medium supplemented with polybrene (4 µg/mL; Sigma-Aldrich, H9268) and Synperonic F108 (100 µg/mL; Sigma-Aldrich, 07579) to enhance transduction efficiency. One hour before infection, cells were dissociated with Accutase and replated at a 1:2.5 split ratio. Cells were exposed to the lentiviral suspension for 3 h, washed twice with DMEM/F-12 and cultured in Essential 8 medium supplemented with RevitaCell.

### CRISPR-LICHT organoid screening workflow

CRISPR-LICHT couples lentiviral delivery of sgRNAs with lineage barcoding, enabling simultaneous identification of the genetic perturbation and reconstruction of clonal expansion dynamics during organoid development. In this study, lineage barcodes were delivered together with sgRNAs in the CRISPR-LICHT lentiviral vector and were used to assign sequencing reads to individual progenitor-derived lineages.

In the original CRISPR-LICHT implementation, unique cell barcodes are additionally introduced during PCR amplification using the UMI-NOPE strategy to reduce amplification bias. In the present study, the UMI-NOPE step was omitted to maximize lineage recovery and effective library coverage. Cell barcode sequences were retained in the amplified region but were not used for downstream filtering or quantification.

For both screens, hESCs were transduced at low multiplicity of infection, corresponding to approximately 15–20% infected cells, to favour single sgRNA integration events. Transduced cells were enriched before organoid generation either by GFP-based FACS sorting or by puromycin selection with 0.75 µg/mL puromycin, according to the experimental condition. GFP-based sorting was performed using a BD FACS Melody cell sorter, with gates defined based on negative and untreated control populations. Differentiation was initiated 24h after enrichment, and eCas9 expression was induced at day 5 by 4-OHT treatment, as described above.

The SOX2-focused library was used to compare two initial enrichment strategies and two endpoint processing strategies. This generated four experimental conditions: GFP enrichment followed by final dTomato sorting, GFP enrichment followed by bulk endpoint harvest, puromycin selection followed by final dTomato sorting, and puromycin selection followed by bulk endpoint harvest. For each condition, 20 organoids were generated and pooled in groups of 10 organoids, yielding two biological replicates per condition (Table S7; Fig. S3).

The Tdark-focused library was screened using the standard FACS-based CRISPR-LICHT workflow, consisting of GFP enrichment before organoid generation and dTomato sorting at the endpoint. A total of 160 organoids were generated and pooled in groups of 6–7 organoids, yielding 25 biological replicates (Table S7; Fig. S4).

Organoids were cultured until day 30–35, corresponding to the SOX2-positive neural progenitor-enriched window selected for screening. Organoid numbers were selected to maintain sufficient library representation and achieve at least 2,000 cells per sgRNA, minimizing dropout and guide underrepresentation.

### Organoid dissociation, endpoint sorting and genomic DNA extraction

At the endpoint, organoids were pooled in groups of 6–7 and dissociated into single-cell suspensions by incubation at 37 °C on a Thermo-Shaker in a 10:1 mixture of Accutase (Sigma-Aldrich, A6954) and 1×? Trypsin (Gibco, 15400). Dissociated cells were filtered through a 35 µm cell strainer before downstream processing.

Depending on the experimental condition, endpoint samples were either sorted by FACS to enrich for dTomato-positive cells, marking activation of the inducible eCas9 cassette, or processed directly as bulk dissociated populations without final sorting. Endpoint dTomato sorting was performed using a BD FACS Melody cell sorter, with gates defined based on negative and untreated control populations (Fig. S3 and Fig. S4). The bulk harvest strategy was used to assess whether bypassing final dTomato sorting preserved sufficient screening information while avoiding an additional FACS step.

Genomic DNA was extracted separately from each pooled sample using a phenol-chloroform extraction protocol. Briefly, samples were mixed with 400 µL of equilibrated phenol (pH 8.0), vigorously shaken for 30 s, and transferred to Phase Lock Gel tubes (Catalog No. 2302820). Following centrifugation at maximum speed for 5 min at room temperature, the upper aqueous phase was collected and subjected to a second phenol extraction, followed by chloroform extraction to remove residual phenol. DNA was precipitated by adding NaCl (0.2 M final concentration), glycogen (0.5 µL), and 2.5–3 volumes of absolute ethanol, followed by overnight incubation on ice. Samples were centrifuged for at least 60 min at 4 °C, and the resulting DNA pellets were washed twice with 70–75% ethanol and once with 70% ethanol. After air-drying for 15–20 min, DNA was resuspended in 0.5× EB buffer and incubated at 55 °C for 1–2 h to facilitate dissolution. DNA concentration was assessed using a NanoDrop spectrophotometer, and the extracted DNA was subsequently used for NGS library preparation.

### PCR amplification and next-generation sequencing

Genomic DNA extracted from dissociated organoid samples was subjected to 12–20 freeze-thaw cycles before amplification. Amplicon libraries were generated by two sequential 50 µL qPCR reactions designed to recover the sgRNA-lineage barcode-containing region and introduce sequencing adaptors and sample-specific barcodes. PCR was performed using up to 50,000 cells per reaction with AmpliTaq Gold 360 Master Mix (Life Technologies, 4398881) and EvaGreen Plus Dye, 20X in water (Biotium, 76423-470).

In the first PCR, target regions were amplified using a mixture of forward primers incorporating staggered random nucleotides to increase sequence diversity (CE1102for, CE1107for, CE1108for, CE1109for and CE1110for) together with a common reverse primer (CEI-0128-rev). Amplification was monitored by qPCR and stopped during the exponential phase to minimize amplification bias. PCR products from matched input samples were pooled and used as template for a second PCR to introduce Illumina sequencing adaptors and sample-specific indices using forward primers from the CE1111–CE1207 series and reverse primer CE735. The second PCR was likewise monitored by qPCR and stopped during the exponential phase. Full primer sequences and cycling conditions are reported in Table S11 and Table S12.

Final libraries were pooled proportionally, size-selected on a 2% agarose gel, gel-purified, and further cleaned by phenol-chloroform extraction followed by ethanol precipitation. Library size and integrity were assessed using an Agilent High Sensitivity D1000 ScreenTape, confirming a predominant amplicon size of approximately 350 bp (Fig. S5).

Sequencing was performed on an Illumina platform using a single-read 100-cycle configuration. The sequencing design included Read 1 for sgRNA and flanking sequences, Index 1 for lineage barcodes and Index 2 for sample barcodes, with a 30% PhiX spike-in to increase sequence diversity (Table S13). Libraries from the Tdark-focused and SOX2- focused screens were pooled and sequenced within a single lane with a capacity of approximately 1.4 billion reads, with read allocation proportional to library complexity. The Tdark-focused library accounted for the majority of sequencing depth, reaching up to approximately 744 million reads for endpoint samples, whereas the SOX2-focused library required lower coverage, ranging from approximately 20 to 192 million reads per sample, consistent with its reduced library complexity (Table S14).

Sequencing was performed on plasmid library controls, initial T0 samples and endpoint T30– 35 samples. Quality-control analysis confirmed high base quality scores across sequencing cycles, consistent read length of approximately 101 bp and no major technical biases, supporting downstream analysis of sgRNA representation and lineage barcode abundance (Table S15).

### Bioinformatic processing of CRISPR-LICHT sequencing data

Sequencing reads from CRISPR-LICHT libraries were processed using Phantombuster v0.19.2. Reads were demultiplexed based on sample-specific barcodes and parsed to identify the guide RNA sequence and the associated lineage barcode. Each read was thereby assigned to a specific sample, gRNA and lineage barcode combination, allowing reconstruction of gRNA-associated clonal lineages within each biological replicate.

The analysis workflow included demultiplexing, barcode error correction, hopping removal and thresholding. Barcode combinations composed exclusively of guanines were excluded from downstream analyses. Samples with incompatible library preparation or poor technical quality were removed prior to final analysis and are reported in the corresponding supplementary table (Table S7). Read counts were normalized to reads per million (RPM) within each sample to account for differences in sequencing depth.

Lineage barcode abundance was used as a proxy for lineage size. For each gRNA, the distribution of associated lineage sizes was compared with the distribution observed for non-targeting control gRNAs, enabling estimation of the effect of each perturbation on clonal expansion during organoid differentiation.

### RPM filtering and gRNA-level statistical analysis

To reduce noise from weakly represented or non-informative lineages, RPM-based lineage filtering was applied before statistical analysis. Multiple RPM thresholds, ranging from 0 to 100, were evaluated to identify a filtering strategy that best separated non-targeting controls from gRNAs targeting genes with known effects on proliferation (Fig. S6; Table S9).

Non-targeting control gRNAs were expected to show approximately uniform P-value distributions, whereas proliferation-associated control gRNAs were expected to show an enrichment of low P-values. An RPM threshold of 40 provided the best separation between these control categories and was therefore used for downstream analyses (Fig. S6). Lineages with fewer than 40 RPM were excluded, and gRNA-level statistics were calculated on the retained lineage distributions.

For each gRNA, log₂ fold change in lineage size was calculated relative to the mean lineage size observed for non-targeting controls. gRNAs with P ≤ 0.01 and negative log₂ fold change were classified as significant negative-effect gRNAs, corresponding to perturbations associated with reduced lineage expansion. Complete gRNA-level results, including P-values, log₂ fold changes, fold changes, number of measured lineages and guide category, are reported in Table S10.

### Gene-level scoring and hit nomination

To identify gene-level hits from gRNA-level data, guide effects were aggregated across the gRNAs targeting each gene. gRNAs associated with fewer than 10 detected lineages were excluded from gene-level scoring to reduce the influence of underrepresented perturbations.

A cumulative scoring system was applied based on gRNA-level statistical significance. Each gRNA was assigned 4 points for P ≤ 0.01, 2 points for P ≤ 0.05 and 1 point for P ≤ 0.10. Guide-level scores were summed for each gene and adjusted for the number of observed gRNAs using a scaling factor of 5/n, where *n* represents the number of gRNAs retained for that gene. This generated a final gene score ranging from 0 to 20.

This scoring strategy prioritized genes supported by concordant effects across multiple independent gRNAs, rather than candidates driven by a single strongly depleted guide.

Genes were ranked according to final gene score and supporting guide-level evidence. Complete gene-level results, including final gene score, number of retained gRNAs, average log₂ fold change and minimum P-value, are reported in Table S10.

### Analysis of the SOX2-focused screen

The SOX2-focused screen was analysed using the same computational framework applied to the Tdark-focused library. Sequencing reads were demultiplexed and parsed to extract sample identifiers, gRNA sequences and lineage barcodes, allowing reconstruction of gRNA-associated lineage-size distributions across the four workflow conditions.

Because the SOX2-focused library did not include an internal set of non-targeting control gRNAs, non-targeting controls from the Tdark-focused screen were used as a shared reference background for normalization and P-value calibration. This enabled direct comparison of gRNA-level effects across the SOX2-focused workflow conditions while maintaining analytical consistency with the Tdark-focused screen.

For each condition, gRNA-level log₂ fold changes, P-values and lineage counts were calculated as described above. Complete results for all SOX2-focused gRNAs across the four tested conditions are reported in Table S8.

### Protein class annotation

Protein class annotation was performed for genes included in the SOX2-focused library, the full Tdark-focused library and the top-ranked Tdark-focused hits. Genes were assigned to broad functional categories based on available protein annotations and manual curation, including transcription factors or chromatin regulators, ion channels, enzymes, signaling proteins, structural or cytoskeletal proteins, scaffold or adaptor proteins, transporters, receptors, membrane-trafficking proteins, mitochondrial proteins and other or uncharacterized proteins (Tables S4 and S5).

For the SOX2-focused library, protein class annotation was used to describe the functional composition of the targeted reference gene set. For the Tdark-focused library, annotation was performed both on the full 120-gene candidate set and on the nominated hit genes to assess whether specific protein classes were represented among genes affecting lineage expansion.

### Expression-informed prioritization of Tdark-focused hits

To prioritize Tdark-focused hits for targeted functional follow-up, genes nominated by the CRISPR-LICHT screen were integrated with expression data from published human brain organoid and developmental resources. For the main prioritization analysis, we used the cerebral organoid dataset from Lancaster et al. (2017), which was generated using a differentiation protocol comparable to that used in this study. Expression values for the 28 nominated Tdark-focused hits were extracted, ranked according to expression level, reported as baseMean, and integrated with the corresponding CRISPR-LICHT gene scores (Fig. 6B).

As an additional in vivo developmental reference, expression data for the 28 nominated hits were extracted from the Allen Brain Atlas Developing Human Brain Tissue Gene Expression Profiles, focusing on selected early cortical and forebrain regions at 8 and 9 post-conception weeks (pcw) (Fig. S7; Table S17).

CDC42EP4 was selected for targeted functional follow-up because it combined a high screen-derived gene score with high expression among the nominated candidates in the Lancaster et al. organoid dataset, expression in early human cortical and forebrain regions in the Allen developing human brain dataset, and functional annotation linking the encoded protein to cytoskeletal organization, actin-associated processes, cell shape regulation and polarity control.

### Organoids electroporation

Cerebral organoids were maintained under antibiotic-free conditions prior to electroporation. Electroporation was performed in 35-days-old cerebral organoids (week 6). During the electroporation, cerebral organoids were placed in a 5 cm dish filled with PBS under a stereomicroscope, and using a glass microcapillary, 1 to 2 ml of plasmid DNA diluted in PBS was injected together with Fast Green (final concentration 0.1%, from Sigma-Aldrich) into different ventricles of the organoids (Buchsbaum et al., 2020; Chinnappa et al., 2022; Tynianskaia et al., 2023).

Plasmid DNA concentration was as follows: non targeting guide in contaminant with pCAGG-GFP at a 3:1 ratio, CDC16 CRISPR guide in contaminant with pCAGG-GFP at a 3:1 ratio and CDC42EP4 CRISPR guide in contaminant with pCAGG-GFP at a 3:1 ratio. Cerebral organoids were subsequently electroporated with five pulses applied at 65 V for 50 ms each at intervals of 500 ms using an ECM 830 electroporator (BTX). Following electroporation, cerebral organoids were incubated for an additional 24 hours in antibiotic-free media, then transferred to normal media. Organoids were kept in culture and fixed 2 days after electroporation.

Cerebral organoids were fixed overnight at 4°C in 4% paraformaldehyde (PFA) prepared in 120 mM phosphate buffer (pH 7.4). Following fixation, organoids were processed for immunofluorescence analysis.

### Immunofluorescence on electroporated organoids and image analysis

Samples were embedded in OCT (Bio-Optica, OCT 05-9801) and frozen using isopentane chilled with dry ice. Cryosections were cut at 20 μm using a Leica CM1950 UV cryostat and stored at −20 °C until use.

Immunofluorescence staining was performed as previously described (Taverna *et al*., 2016). Briefly, slides containing cryosections were dried for 20 min at 37 °C and briefly washed with PBS before antigen retrieval was performed in 10 mM citrate buffer, pH 6.0, for 1 h at 70 °C. Following a 5 min PBS wash, slides were incubated with blocking buffer for 30 min at room temperature. Sections were then incubated overnight at 4 °C with primary antibodies diluted in blocking buffer. The following primary antibodies were used: goat anti-SOX2 (R&D Systems, AF2018, 1:200), rabbit anti-TBR2 (Abcam, ab261913, 1:250) and rabbit anti-NEUN (Invitrogen, PA5-78499, 1:250).

The following day, slides were washed three times for 5 min with PBS before incubation with secondary antibodies, diluted 1:1000 in blocking buffer together with DAPI. Slides were incubated for 1–2 h at room temperature, washed three times for 5 min in PBS and mounted with Mowiol. After Mowiol solidification, slides were stored at 4 °C before imaging. Images were acquired using a Zeiss LSM980-NLO confocal microscope with a 40× oil objective.

For cortical organoids, only ventricles facing the organoid surface were included in the analysis, as centrally located ventricles may be affected by limited oxygen and nutrient diffusion. Quantification was performed using Fiji (Schindelin *et al*., 2012), data processing was conducted in Microsoft Excel, and data visualization was performed using GraphPad Prism. All image analyses were performed manually on single optical planes. Cortical regions, including the ventricular zone (VZ) and subventricular zone (SVZ), were manually delineated based on their characteristic cytoarchitecture observed by DAPI staining. The VZ was identified as the densely packed layer of elongated, radially oriented nuclei surrounding the ventricular lumen, whereas the SVZ was defined as the adjacent basal region containing more loosely arranged nuclei. The accuracy of these boundaries was confirmed using SOX2 immunostaining, based on the distribution and characteristic radial orientation of SOX2- positive progenitor nuclei around the ventricle.

