## Supplementary material for "An improved CRISPR-LICHT screen of 120 “Tdark” genes illuminates new biologically relevant SOX2 targets in human brain organoids": Pozzolini et al Figures S1-S8

### 6. Supplementary

#### Supplementary Figures

##### Figure S1. Adapted Lancaster-based protocol used for brain organoid generation and representative quality-control checkpoints.

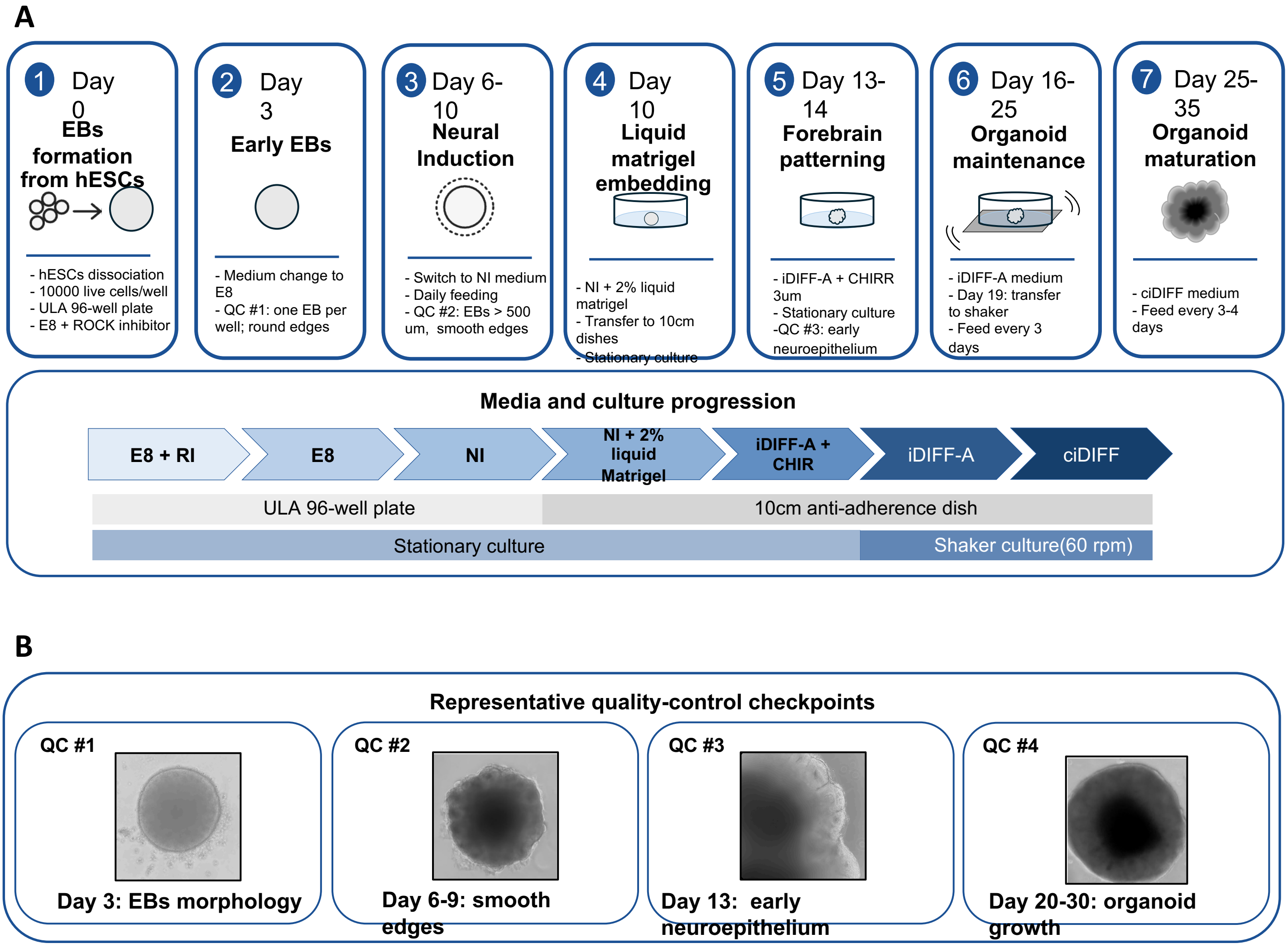


**Figure S1. Adapted brain organoid differentiation protocol and representative quality-control checkpoints.**

**(A)** Schematic overview of the adapted Lancaster-based protocol used for generation of brain organoids from hESCs. Embryoid bodies (EBs) were formed in ultra-low-attachment 96-well plates and subsequently progressed through neural induction, liquid Matrigel embedding, forebrain patterning and later organoid maturation. The scheme indicates the main culture stages, medium changes and transition from stationary to shaker culture. For screening experiments, 4-hydroxytamoxifen (4-OHT) was added at day 5 to induce eCas9 expression.

**(B)** Representative morphological quality-control checkpoints during organoid differentiation. Representative images show EB morphology at day 3, smooth-edged EBs at day 6, early neuroepithelium at day 13 and organoid growth at later stages (day 20–30). These criteria were used to monitor differentiation quality throughout organoid generation.

RI, ROCK inhibitor; E8, Essential 8 medium; NI, neural induction medium; iDIFF-A, induction differentiation medium A; cDIFF, cerebral differentiation medium.

**Figure S2. Lentiviral CRISPR-LICHT backbone used for sgRNA and lineage barcode library generation.**

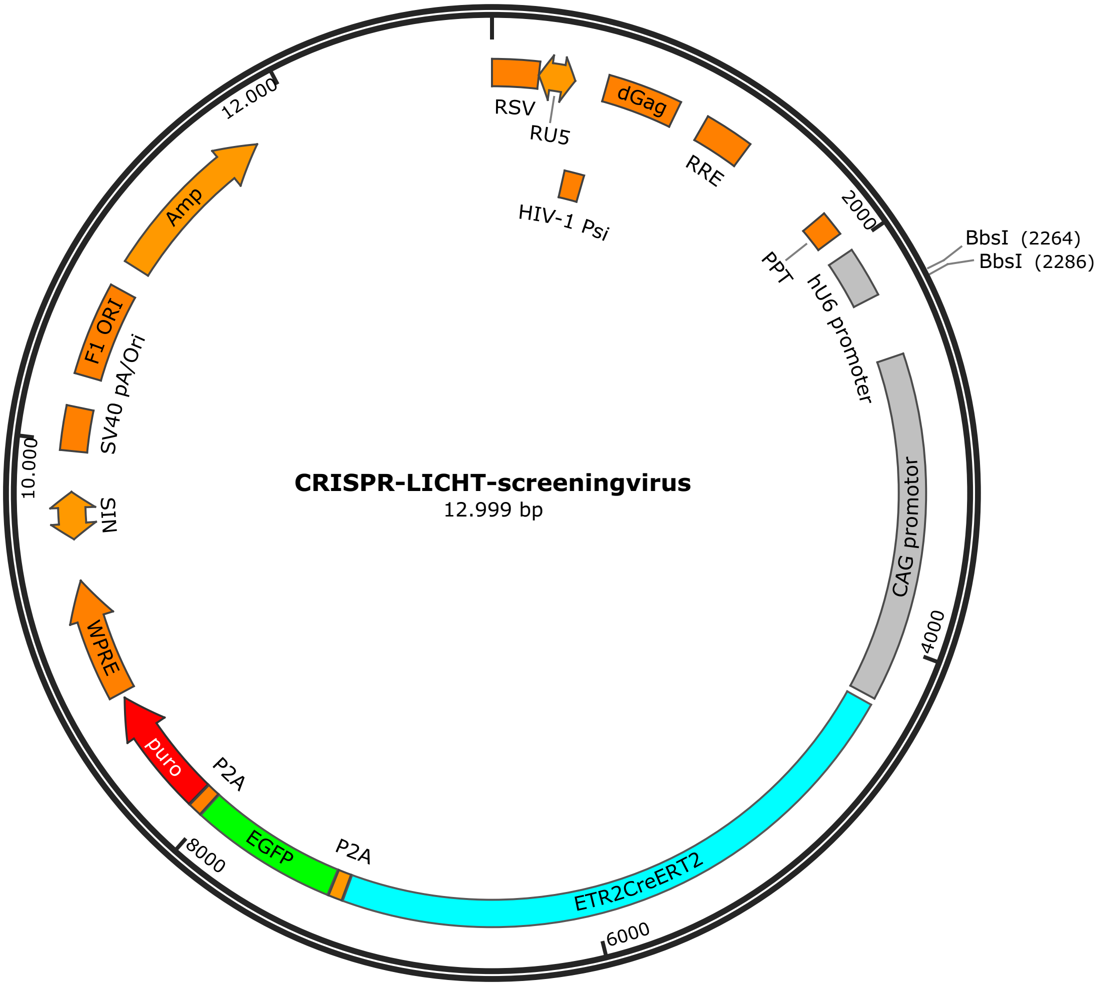


**Figure S2. Lentiviral CRISPR-LICHT backbone used for sgRNA and lineage barcode library generation.**
Schematic representation of the human screening CRISPR-LICHT lentiviral backbone used for cloning of the SOX2-focused and Tdark-focused sgRNA libraries. The vector contains the U6-driven sgRNA cassette, the lineage barcode-containing region used for clonal tracking, and a CAG-driven selection cassette encoding CreERT2, EGFP and puromycin resistance. In the inducible CRISPR-LICHT system, CreERT2-mediated recombination activates eCas9 expression following 4-hydroxytamoxifen treatment in the recipient hESC line. Major vector elements are indicated.

**Figure S3. Effect of initial enrichment strategy on reporter-defined cell populations in SOX2-focused organoids.**

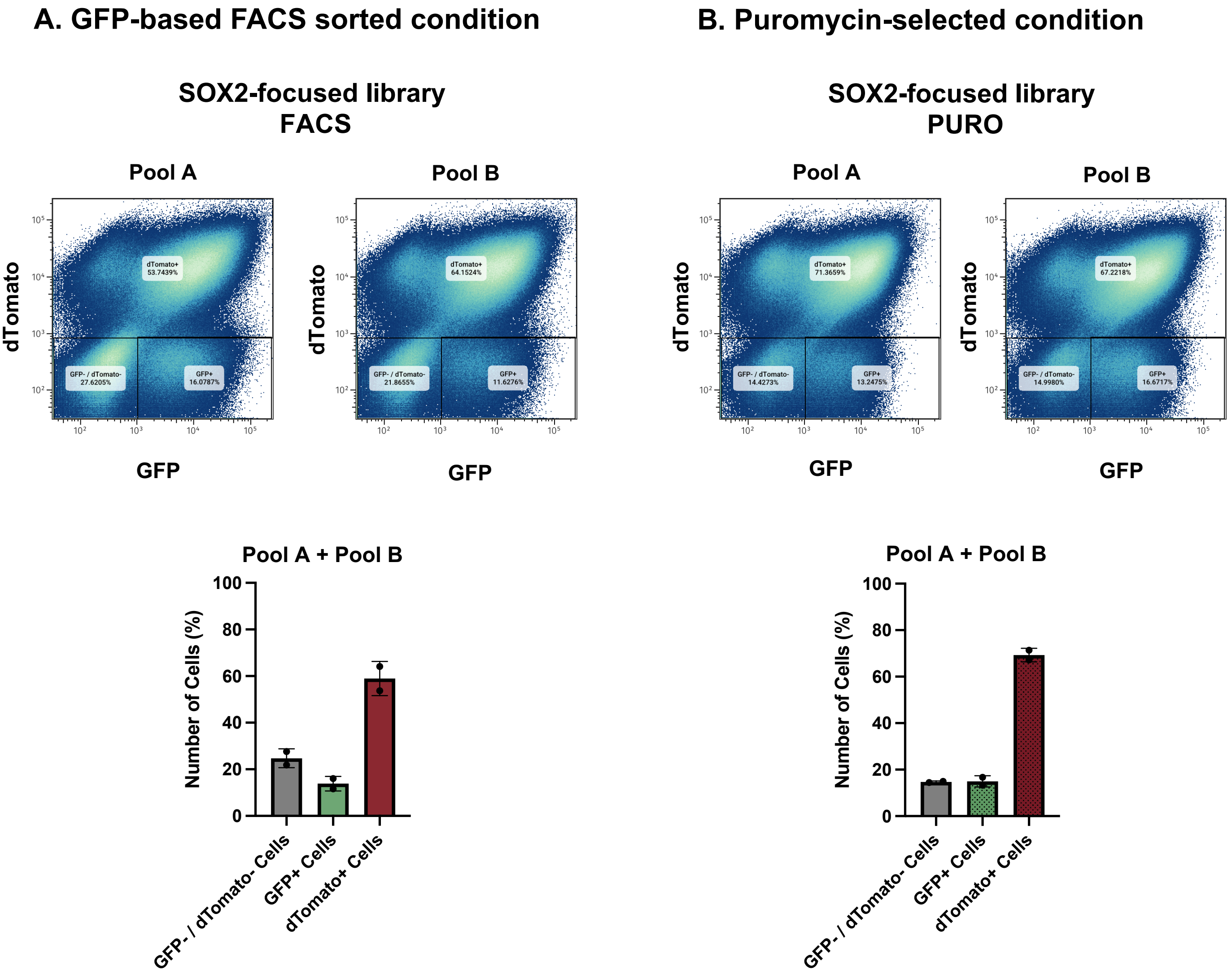


**Figure S3. Effect of initial enrichment strategy on reporter-defined cell populations in SOX2-focused organoids.**
**(A,B)** Flow cytometry analysis of day 30 SOX2-focused organoids generated after enrichment of transduced hESCs by GFP-based FACS sorting **(A)** or puromycin selection **(B)**. Representative GFP/dTomato plots are shown for two independent organoid pools per condition, together with quantification of reporter-defined populations. Puromycin selection increased the proportion of dTomato-positive cells and reduced the GFP-negative/dTomato-negative fraction compared with GFP-based enrichment. Gates were defined using negative and untreated control populations. Each point represents one pool of 10 organoids; bars show mean ± s.e.m.

**Figure S4. Endpoint dTomato sorting quality control for the Tdark-focused CRISPR-LICHT screen**

#### A. FACS analysis of endpoint Tdark-focused organoid samples

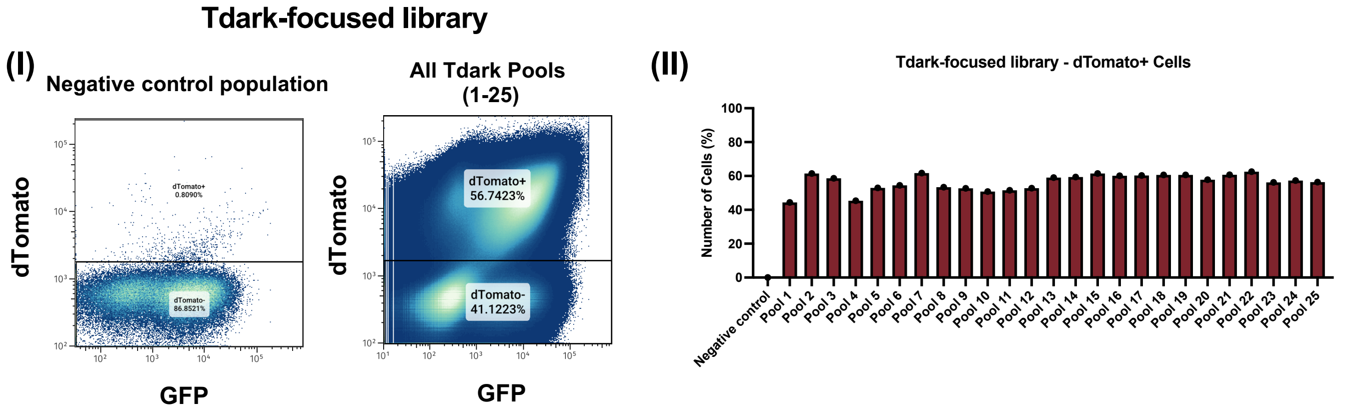


#### B. Number of dTomato-sorted cells recovered from each Tdark-focused organoid pool

| Tdark-organoids pool | N. Sorted cells |
|---|---|
| Pool 1 | 1x10^6 cells |
| Pool 2 | 600000 cells |
| Pool 3 | 770000 cells |
| Pool 4 | 1x10^6 cells |
| Pool 5 | 1x10^6 cells |
| Pool 6 | 1,1x10^6 cells |
| Pool 7 | 855000 cells |
| Pool 8 | 1,1x10^6 cells |
| Pool 9 | 1,2x10^6 cells |
| Pool 10 | 1,1x10^6 cells |
| Pool 11 | 1,1x10^6 cells |
| Pool 12 | 1x10^6 cells |
| Pool 13 | 600000 cells |
| Pool 14 | 625000 cells |
| Pool 15 | 830000 cells |
| Pool 16 | 900000 cells |
| Pool 17 | 750000 cells |
| Pool 18 | 1,1x10^6 cells |
| Pool 19 | 1,1x10^6 cells |
| Pool 20 | 960000 cells |
| Pool 21 | 640000 cells |
| Pool 22 | 755000 cells |
| Pool 23 | excluded* |
| Pool 24 | 1x10^6 cells |
| Pool 25 | 770000 cells |

*Pool 23 was excluded from downstream processing because of a technical issue after cell sorting

**Figure S4. Endpoint dTomato sorting quality control for the Tdark-focused CRISPR-LICHT screen.**

**(A)** Flow cytometry analysis of endpoint Tdark-focused organoid samples before dTomato-positive cell sorting. Representative GFP/dTomato plots are shown for the negative control population and pooled Tdark-focused samples (I), together with quantification of the dTomato-positive fraction across the 25 organoid pools prepared for endpoint sorting (II). Each pool contained 6–7 organoids. Gates were defined using negative and untreated control populations.

**(B)** Number of dTomato-sorted cells recovered from each Tdark-focused organoid pool. Sorted cells from pool 23 were not included in downstream sequencing analysis because of a technical issue after collection.

#### Figure S5. Quality control of CRISPR-LICHT sequencing libraries and structure of the sequenced amplicon

##### A. Amplicon-library quality control

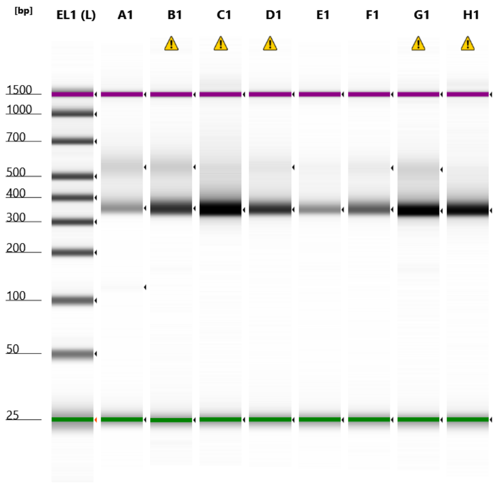


**Sample Info**

| Well | Conc. [pg/µl] | Sample Description |
|---|---|---|
| EL1 | 2350 | Electronic Ladder |
| A1 | 666 | T35 TDARK |
| B1 | 1320 | T0 TDARK |
| C1 | 2220 | plasmid TDARK |
| D1 | 1030 | T35 SOX2-SORTED |
| E1 | 391 | T0 SOX2-SORTED |
| F1 | 824 | T35 SOX2-PURO |
| G1 | 2300 | T0 SOX2-PURO |
| H1 | 1550 | plasmid SOX2 |

##### B. Structure of the CRISPR-LICHT amplicon

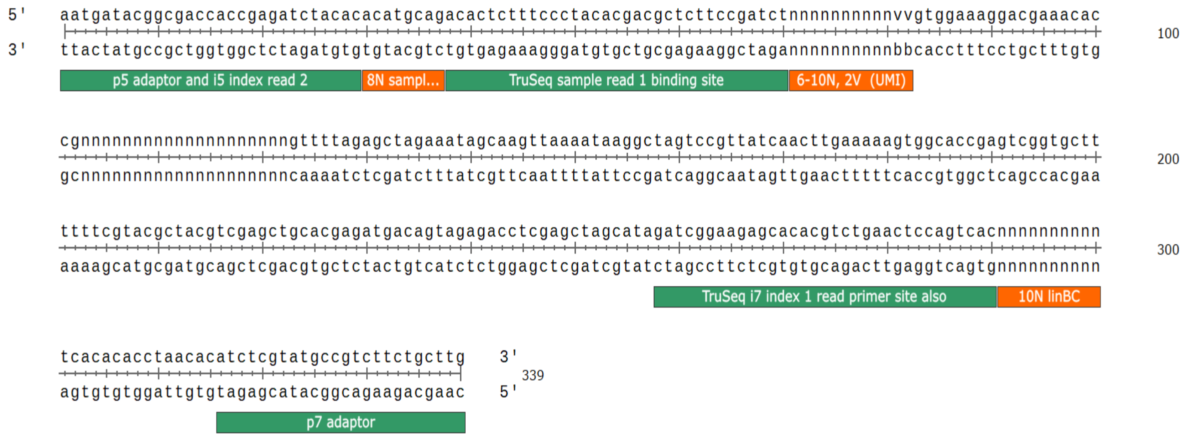


**Figure S5. Quality control of CRISPR-LICHT sequencing libraries and structure of the sequenced amplicon.**

**(A)** Representative quality-control analysis of amplified CRISPR-LICHT sequencing libraries generated from plasmid controls, T0 samples and endpoint samples from the Tdark-focused and SOX2-focused screens. Electrophoretic profiles show successful amplification of sequencing-compatible amplicons of the expected size across representative samples. **(B)** Schematic representation of the sequenced CRISPR-LICHT amplicon, showing the relative position of sequencing adaptor regions, sample barcode, sgRNA-containing region, UMI/NOPE-related sequence elements and lineage barcode regions used for clonal lineage assignment.

**Figure S6. Library representation, lineage recovery and RPM threshold selection for the Tdark-focused screen.**

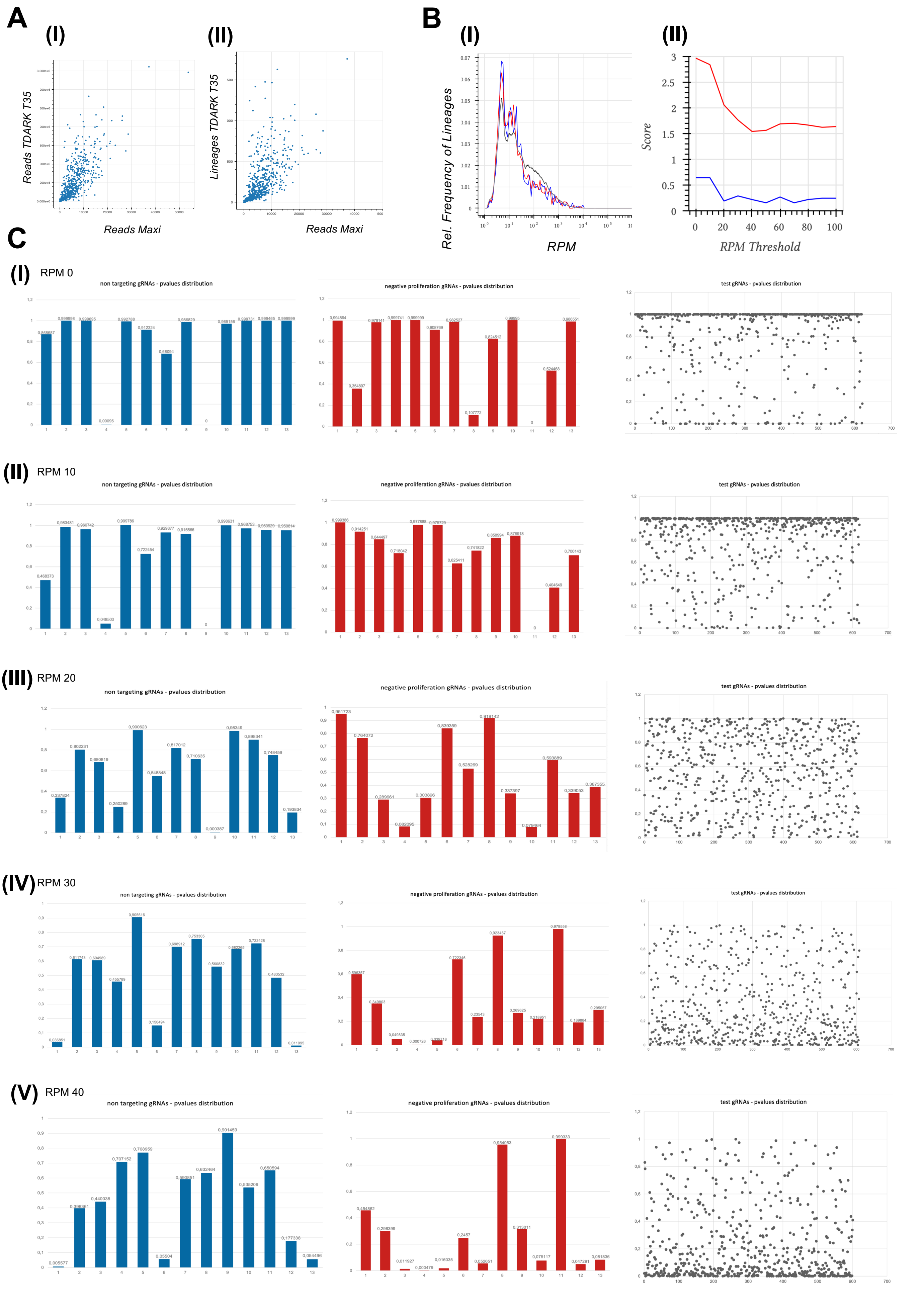

**Figure S6. Library representation, lineage recovery and RPM threshold selection for the Tdark-focused screen.**
**(A)** Correlation analyses comparing plasmid-library representation with endpoint recovery in the Tdark-focused screen. Scatter plots show the relationship between plasmid-library read abundance and endpoint read abundance, and between plasmid-library read abundance and the number of detected endpoint lineages.
**(B)** Distribution of lineage RPM values and scoring of candidate RPM thresholds. RPM thresholds from 0 to 100 were evaluated to identify a filtering strategy that retained informative lineages while reducing noise from weakly represented clones. The selected threshold of 40 RPM is indicated.
**(C)** P-value distributions for non-targeting control gRNAs, proliferation-associated control gRNAs and test gRNAs across increasing RPM thresholds. Non-targeting controls were expected to show approximately uniform P-value distributions, whereas proliferation-associated controls were expected to show enrichment of low P-values. An RPM threshold of 40 provided improved separation between these control categories and was used for downstream gRNA-level analysis.

**Figure S7. Expression of the 28 nominated Tdark hit genes across early human cortical and forebrain regions in the Allen Human Brain Atlas.**

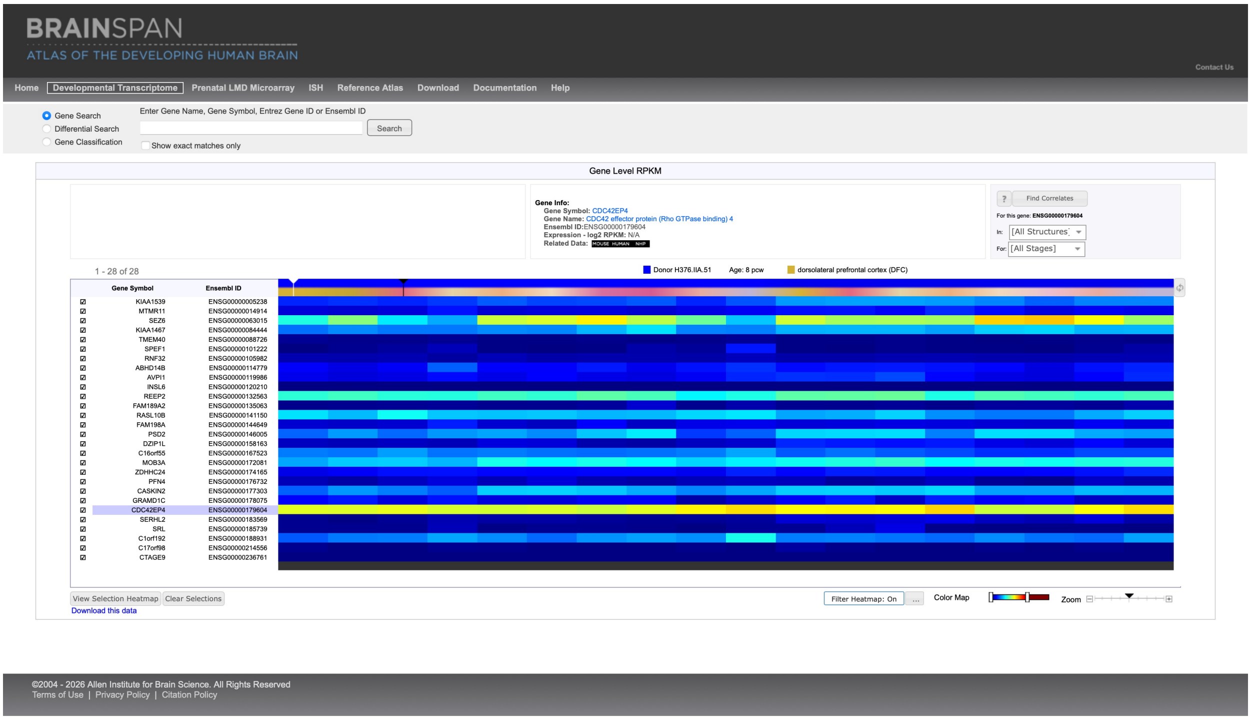


**Figure S7. Expression of the 28 nominated Tdark hit genes across early human cortical and forebrain regions in the Allen Human Brain Atlas.**

Heatmap showing expression levels of the 28 top-ranked candidate genes identified in the Tdark-focused CRISPR-LICHT screen across selected cortical and forebrain regions of the developing human brain at 8 and 9 post-conception weeks (pcw), based on data from the Allen Human Brain Atlas. Regions include primarily cortical and forebrain areas, with neocortical subdivisions such as frontal, parietal, temporal and occipital cortex where annotated. The heatmap was used to assess whether nominated hits are expressed in early human brain regions relevant to cortical and forebrain development. Specific expression values and region-level annotations are reported in Table S17.

**Figure S8. Progenitor marker analysis and criteria for spatial and apoptotic cell classification following organoid electroporation**

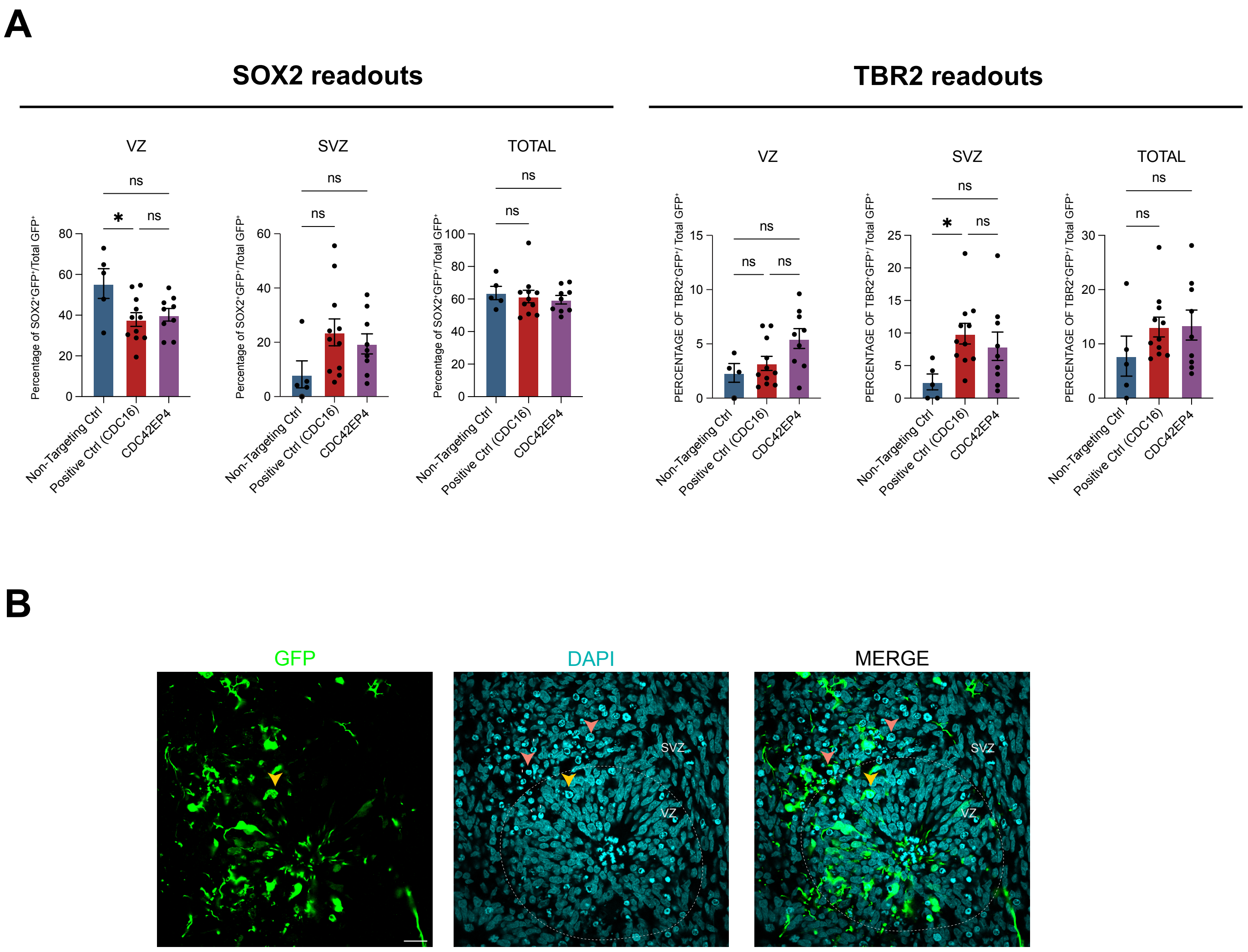


**Figure S8. Progenitor marker analysis and criteria for spatial and apoptotic cell classification following organoid electroporation**

**(A)** Quantification of $GFP^+SOX2^+$ (left) and $GFP^+TBR2^+$ (rigth) cells within the VZ, SVZ, and total ventricular region. Graphs show the percentage of GFP-positive cells in the indicated regions, relative to the total number of GFP-positive cells, that are double-positive for the different stem markers. Data are presented as mean ± SEM. Each dot represents an individual ventricle analyzed. A total of 5–11 electroporated ventricles from 6-7 independent organoids from two independent batches were analyzed per condition. Statistical significance was determined using one-way ANOVA with Turkey's multiple comparisons test. ns, not significant; $*P < 0.05$; $**P < 0.01$.

**(B)** Apoptotic cells were identified by pyknotic nuclei showing condensed or fragmented DAPI staining, consistent with apoptotic morphology. Pyknotic nuclei were manually counted within a 100-µm-wide region of the GFP-electroporated area, and the total number of GFP+ apoptotic cells was quantified for each section. The orange arrowhead indicates a pyknotic electroporated GFP+ cell. Pink arrowheads indicate pyknotic non-electroporated GFP- cells. VZ-like and SVZ-like regions were identified using DAPI staining to highlights nuclei. VZ cells show a characteristic arrangement of elongated nuclei radially aligned around the central lumen, consistent with ventricular zone architecture. SVZ-like regions were identified by first delineating ventricular rosettes using DAPI staining. Regions of interest (ROIs) corresponding to the SVZ-like area were identified as immediately adjacent to the ventricular lumen based on the characteristic cytoarchitecture of the rosette. Scale bar, 20 µm.
