## Supplementary material for "An improved CRISPR-LICHT screen of 120 “Tdark” genes illuminates new biologically relevant SOX2 targets in human brain organoids": Pozzolini et al Tables S1-S17: Table S1, S11-S14 copia.pdf

### Supplementary Tables

#### Table S1. Pooled genetic screens in human brain organoids/assembloids and coverage of SOX2

Overview of major pooled perturbation studies performed in human brain organoids or forebrain assembloids. The table indicates whether SOX2 itself was perturbed and whether a module of direct SOX2 targets was screened. “SOX2 targets tested systematically” denotes an a priori, curated set of direct SOX2 target genes. To our knowledge, none of the listed pooled screens included SOX2 or a dedicated SOX2-target module in human cortical organoids.

| Study | Year | Model | Screen modality | Gene set focus | SOX2 perturbed? | SOX2 targets tested systematically? | Key note | Reference (journal; DOI) |
|---|---|---|---|---|---|---|---|---|
| Esk et al., 2020 | 2020 | Human cerebral organoids | CRISPR-LICHT pooled LOF; dual lineage/cell barcodes | 173 microcephaly candidate genes | No | No | Introduces lineage-resolved pooled screening; identifies IER3IP1 as ER-secretion regulator linked to brain size | https://doi.org/10.1126/science.abb5390 |
| Li et al., 2023 (CHOOSE) | 2023 | Human cerebral organoids | Pooled LOF with single-cell RNA-seq readout | 36 high-risk ASD genes (transcriptional regulators) | No | No | Single-cell, lineage-aware barcoding with inducible eCas9; cell type–resolved phenotypes | https://doi.org/10.1038/s41586-023-06473-y |
| Meng et al., 2023 | 2023 | Human forebrain assembloids | Two pooled CRISPR screens (generation and migration) | 425 neurodevelopmental disorder genes | No | No | Maps NDD genes to interneuron generation/migration in assembloids | https://doi.org/10.1038/s41586-023-06564-w |
| Bury et al., 2024 | 2024 | Human cortical organoids | Lineage tracing (COR-LT); not a pooled CRISPR screen | — | Not applicable | No | Records progenitor lineages and neuronal production dynamics | https://doi.org/10.1016/j.celrep.2024.114862 |
| Hendriks et al., 2024 | 2024 | Human fetal brain organoids (FeBOs) | CRISPR-mutated clonal organoids (not pooled) | Method platform; cancer modeling use case | No | No | Long-term expanding fetal tissue–derived organoids; CRISPR amenable | https://doi.org/10.1016/j.cell.2023.12.012 |

**Table S11. Primers used for PCR amplification and library preparation**

Sequences are provided in 5′–3′ orientation. Inner PCR primers include variable-length random nucleotides to increase sequence diversity. Outer PCR primers introduce sequencing adapters and sample indices.

| Primer name | Sequence (5'–3') | Function |
|---|---|---|
| CE1102for | CCCTACACGACGCTCTTCCGATCT(N)xVVGTGGAAAGG ACGAAACACCG | Inner PCR forward (staggered) |
| CE1107for | same backbone with variable N | Inner PCR forward (staggered) |
| CE1108for | same backbone with variable N | Inner PCR forward (staggered) |
| CE1109for | same backbone with variable N | Inner PCR forward (staggered) |
| CE1110for | same backbone with variable N | Inner PCR forward (staggered) |
| CEI-0128-rev | CGACGTTTAAACTGCTATCCTGCAG | Inner PCR reverse |
| CE1111–CE1207 | AATGATACGGCGACCACCGAGATCTACAC(index)ACAC TCTTTCCCTACACGACGCTCTTCCGATCT | Outer PCR forward (indexed) |
| CE735 | CAAGCAGAAGACGGCATACGAGATGTGTTAGGTGTG | Outer PCR reverse |

(N)x indicates 6–10 random nucleotides introduced to increase sequence diversity.

**Table S12. PCR cycling conditions used for library amplification**

Cycling parameters for PCR1 and PCR2 steps, including temperatures, durations and cycle numbers. Amplification cycles were monitored and terminated during the exponential phase to minimise amplification bias.

| Step | Temperature | Time | Cycles | Notes |
|---|---|---|---|---|
| **PCR1** – Initial denaturation | 95°C | 5 min | 1 | |
| **PCR1** – Pre-amplification | 95°C / 57°C / 72°C | 30 s / 25 s / 45 s | 5 | |
| **PCR1** – qPCR cycles | 95°C / 57°C / 72°C | 25 s / 25 s / 30 s | variable | Monitored, stopped in exponential phase |
| **PCR1** – Final extension | 72°C | 15 s | 1 | |
| **PCR2** – Initial denaturation | 95°C | 5 min | 1 | |
| **PCR2** – Pre-amplification | 95°C / 65°C / 72°C | 25 s / 25 s / 45 s | 3 | |
| **PCR2** – qPCR cycles | 95°C / 72°C | 25 s / 60 s | variable | Monitored, stopped in exponential phase |
| **PCR2** – Final extension | 72°C | 15 s | 1 | |

**Table S13. Sequencing configuration and library design**

Summary of sequencing platform, read structure and library features, including read allocation for sgRNA sequences, lineage barcodes and sample indices.

| Parameter | Description |
|---|---|
| Platform | Illumina sequencing |
| Read configuration | Single-read, 100 cycles |
| Read 1 | sgRNA and flanking sequence |
| Index 1 (i7) | Lineage barcode (LB) |
| Index 2 (i5) | Sample barcode |
| PhiX spike-in | 30% |
| Libraries sequenced | TDP (Tdark screen), SP (SOX2-focused screen) |
| Library type | Amplicon sequencing |
| Expected amplicon size | ~350 bp |

**Table S14. Sequencing depth and read allocation across samples**

Number of reads assigned to each sample, grouped by screening condition and experimental timepoint. Read allocation reflects library complexity and experimental design.

| Sample | Screen | Condition | Required reads | % of pool |
|---|---|---|---|---|
| T35 TDARK | Tdark | Endpoint | 744M | 87% |
| T0 TDARK | Tdark | Initial | 60M | 7% |
| Plasmid TDARK | Tdark | Control | 40M | 4.7% |
| T35 SOX2-SORTED | SOX2 | Endpoint | 184M | 33.4% |
| T0 SOX2-SORTED | SOX2 | Initial | 90M | 16.3% |
| T35 SOX2-PURO | SOX2 | Endpoint | 192M | 35% |
| T0 SOX2-PURO | SOX2 | Initial | 90M | 16.3% |

| Sample | Screen | Condition | Required reads | % of pool |
| --- | --- | --- | --- | --- |
| Plasmid SOX2 | SOX2 | Control | 20M | 3.6% |
