## Supplementary material for "An improved CRISPR-LICHT screen of 120 “Tdark” genes illuminates new biologically relevant SOX2 targets in human brain organoids": Pozzolini et al Tables S1-S17: Table S2.pdf

**Table S2. Summary of selected Sox2 target genes: function in mouse and human, disease links, and supporting evidence**

| Gene | Mouse | Human | Key Disease Link | References |
|---|---|---|---|---|
| **ZEB2** | Required for neural crest migration and glial differentiation; loss causes severe CNS malformations. | Transcription factor essential for neuroectodermal differentiation; haploinsufficiency causes Mowat–Wilson syndrome. | Mowat–Wilson syndrome (intellectual disability, epilepsy, brain malformations). | Garavelli et al., 2017 |
| **KCNA2 (Kv1.2)** | Regulates axonal excitability and node of Ranvier conduction; knockout alters neural activity and sleep. | Voltage-gated $K^+$ channel subunit; mutations cause developmental and epileptic encephalopathy (DEE32). | Developmental and epileptic encephalopathy, episodic ataxia. | Syrbe et al., 2015 |
| **GJA1 (Connexin-43)** | Major astrocytic gap-junction protein mediating intercellular communication. | Forms gap junctions and hemichannels in glia; maintains metabolic coupling and astrocyte-neuron signaling. | Oculodentodigital dysplasia (ODDD), arrhythmogenic cardiomyopathy. | Paznekas et al., 2003 |
| **FBRSL1** | Conserved paralog of AUTS2; functionally uncharacterized in mouse. | Nuclear protein implicated in chromatin regulation; mutations cause neurodevelopmental disorder with microcephaly and skeletal defects. | FBRSL1-associated neurodevelopmental disorder. | Kastens et al., 2025 |
| **EGR1 (NGFI-A)** | Immediate early gene induced by neuronal activity; required for memory consolidation and synaptic plasticity. | Activity-dependent transcription factor regulating learning, memory, and stress response. | Implicated in schizophrenia and cognitive disorders. | Pagin et al., 2021 |
| **GRIA2 (GluA2)** | Determines AMPA receptor $Ca^{2+}$ permeability via Q/R RNA editing; critical for neuron survival and plasticity. | Regulates AMPA receptor ion selectivity and synaptic stability; altered in excitotoxicity and epilepsy. | Amyotrophic lateral sclerosis (ALS), epilepsy, cognitive decline. | Efthymiou et al., 2024 |
| **PPP2R2B (PP2A B55β)** | Subunit of PP2A holoenzyme; expressed in cerebellum, involved in neuronal signaling. | Regulatory B55β subunit of PP2A; expanded CAG repeats cause SCA12. | Spinocerebellar ataxia type 12 (SCA12). | Holmes et al., 1999 |
| **EGR2 (KROX-20)** | Required for Schwann cell myelination; knockout causes severe PNS demyelination. | Transcription factor for myelin genes; mutations cause CMT1D and Dejerine–Sottas neuropathy. | Charcot–Marie–Tooth disease, Dejerine–Sottas syndrome. | Pagin et al., 2021 |
| **CNTN1 (Contactin-1)** | GPI-linked adhesion molecule for axonal guidance and myelination. | Mediates neuron–glia interactions and node of Ranvier organization. | Congenital myasthenic syndromes, autoimmune neuropathies. | Labasque et al., 2014 |

| Gene | Mouse | Human | Key Disease Link | References |
|---|---|---|---|---|
| **GADD45G** | Stress-inducible gene regulating cell cycle arrest and neurogenesis; influences neuronal differentiation. | Mediates MAPK signaling and neurite growth via p38–CDC25B axis. | Cancer susceptibility, neurogenesis defects. | Patel et al., 2022 |
| **KCNK2 (TREK-1)** | Two-pore $K^+$ channel modulating excitability, cognition, and neuroprotection. | Mechanosensitive and lipid-gated $K^+$ channel regulating mood and ischemic tolerance. | Depression, ischemia, epilepsy. | Heurteaux et al., 2006 |
| **ARX** | X-linked homeobox TF controlling interneuron migration; knockout causes forebrain patterning defects. | Regulates cortical interneuron specification; mutations lead to epileptic encephalopathy and intellectual disability. | X-linked lissencephaly, Ohtahara syndrome, intellectual disability. | Nieto-Estevez et al., 2024 |
